# Cell-type specific effects of somatostatin on corticocortical information processing in the anterior cingulate cortex

**DOI:** 10.1101/2022.08.02.502518

**Authors:** Therese Riedemann, Bernd Sutor

## Abstract

Inhibitory modulation of glutamatergic information processing is a prerequisite for proper network function. Among the many groups of interneurons, somatostatin-expressing interneurons (SOM-INs) seem to play an important role in the maintenance of physiological brain activity. We have previously shown that bath application of somatostatin (SOM) causes a reduction in pyramidal cell (PC) excitability. However, the mechanisms of action of the peptide on cortical synaptic circuits are still unclear. To gain further insight into the function of SOM-INs, we analyzed the effect of SOM on synaptic transmission in PCs, in SOM-INs and in layer 1 interneurons (L1-INs) of the anterior cingulate cortex. We found that SOM produced pronounced postsynaptic effects in PCs while having little or no postsynaptic effects on either IN type. Accordingly, it is proposed here, that SOM - by acting differentially on SOM-INs and L1-INs - increases spontaneous GABAergic transmission, while at the same time, decreasing spontaneous glutamatergic transmission. In addition, we show that SOM specifically modulates GABA_B_ receptor (GABA_B_R)-mediated synaptic transmission but has little effect on GABA_A_ receptor-mediated (GABA_A_R) transmission.

## 1. Introduction

Within the cerebral cortex, the neuropeptide somatostatin (SOM) is predominantly expressed by a distinct class of GABAergic interneurons (INs) termed somatostatin-positive INs (SOM-INs). SOM is thought to be released upon burst-firing or high frequency action potential discharge of the respective presynaptic neuron [1–3]. The biological actions of the peptide are conveyed via activation of five distinct somatostatin receptor (SSTR) subtypes, SSTR1 to SSTR5, all of which are G-protein coupled receptors [4, 5]. In the postsynaptic target cell, binding of the ligand to SSTRs activates inwardly-rectifying potassium conductances (GIRK) through a direct interaction of the Gβγ subunit of the receptor with the corresponding channels. As a consequence, the postsynaptic cell becomes hyperpolarized and its excitability is reduced. In the brain, actions of SOM are mostly mediated via activation of SSTR2 [6–8]. In agreement with that, autoradiography assays, in situ hybridization studies and single-cell sequencing studies identified SSTR2 as being prominently expressed in the cerebral cortex or as being expressed by the majority of cortical PCs [9–11]. In addition, SSTRs are expressed by different types of GABAergic INs [12].

Cortical GABAergic INs can be classified into molecularly, morphologically and electrophysiologically distinct groups: Parvalbumin-positive INs (PV-INs) represent the largest group and account for around 40% of the overall population of cortical GABAergic INs. SOM-INs in turn represent around 30% of the total IN population. Lastly, another group of INs expresses the 5HT receptor 3A and accounts for around 30% of all GABAergic INs. The latter group is further subdivided into INs either expressing vasoactive intestinal peptide (VIP-INs) or Reelin, but not VIP (non-VIP INs resp. Reelin-INs) [13, 14] and is mostly located in the superficial cortical layers. All IN subtypes inhibit cortical projection neurons in a cell-type specific manner [13, 15–20]. In addition, the majority of GABAergic INs inhibit each other [13, 14].

Several studies have elucidated the role of SOM-INs on sensory stimulus representation and/or its adaption in primary sensory cortex areas such as the somatosensory [21, 22], the visual [23, 24] and the auditory [25–27] cortex. It is likely, that SOM-INs, by endogenous release of the peptide, not only modulate PC activity but also that of other GABAergic INs, thereby increasing the complexity of SOM’s effect on cortical circuits. Indeed, a recent study showed that SOM modulates the synaptic circuitry between cortical PCs and PV-INs by depressing excitatory synaptic input onto PV-INs [28]. However, to date the effect of SOM on SOM-INs themselves and on layer 1 INs (L1-INs) is not known. It remains unclear to which extent endogenously released SOM contributes to the observed effects and how SOM modulates the underlying neuronal circuitry. In order to understand SOM-IN function and their ability to modulate information processing within a given neuronal circuit, understanding of the actions of SOM on synaptic transmission and its modulation of information flow is required. Previous studies have shown that SOM not only depresses presynaptic glutamate [29, 30] but also GABA [31] release in non-cortical neurons, however, its effect on synaptic transmission in cortical neurons remains unknown. Therefore, we analyzed the effects of SOM on glutamatergic and GABAergic synaptic transmission in layer 2/3 PCs (L2/3 PCs), SOM-INs and L1-INs of the anterior cingulate cortex (aCC) and determined its effects on the excitability of the different cell types. Finally, we demonstrate that SOM modulates dendro-somatic integration of feedforward and feedback inputs onto PCs by specifically impacting on GABA_B_R-mediated synaptic transmission in apical dendrites of L2/3 PCs, resulting in altered correlated neuronal activity between L2/3 PCs, L2/3 SOM-INs and L1-INs.

## 2. Results

### 2.1 SSTR and GABA_B_R signaling pathways converge on GIRK channel activation in cortical pyramidal cells

We and others have previously shown that SOM leads to a reduction in excitability of projection neurons [8, 30, 32]. In order to determine whether this effect was conveyed in an activity-dependent manner or not and to determine which somatostatin receptor (SSTR) mediates this effect in the aCC, we initially performed voltage-clamp recordings to detect SOM-induced changes in the holding current. Therefore, we clamped cortical pyramidal cells (PCs) at a holding potential of −60 mV and added SOM to the bath after a baseline recording lasting for 5-10 min. In agreement with previous data by us and others [7, 8, 32], we found that SOM induced an outward current in PCs that could not be blocked by preincubation with TTX, indicating a direct, postsynaptic effect in PCs. Preincubation of cells either with the pan SSTR blocker Cyclosomatostatin or with the specific SSTR2 blocker CYN154806 significantly reduced the SOM-induced outward current. Interestingly though, Cylcosomatostatin proved less effective in blocking the SOM-mediated outward current compared to CYN154806 (Fig 1 A, B).

**Fig 1.**
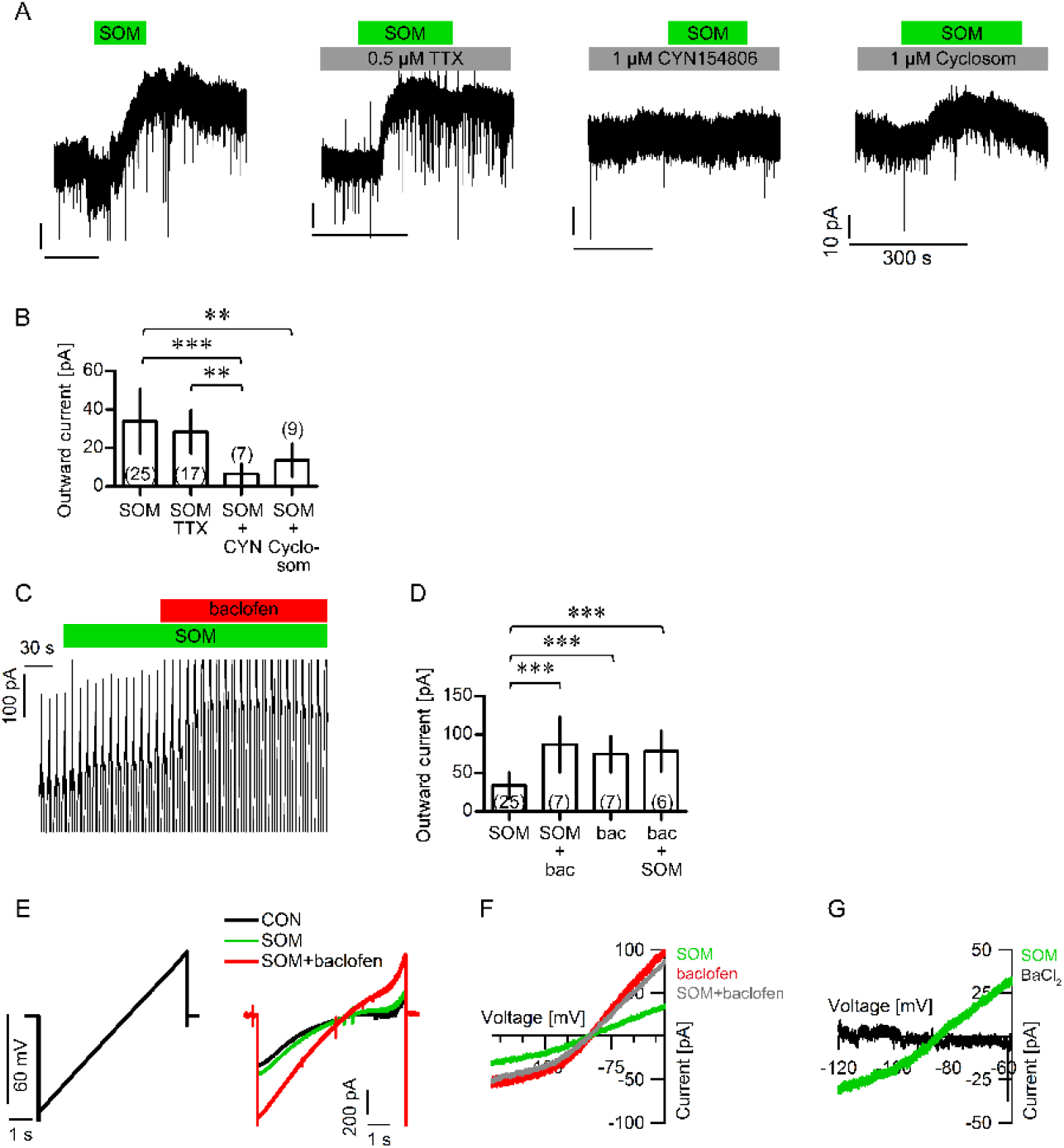
SOM induces GIRK channel activation in L2/3 PCs. A) Representative current traces of L2/3 PCs either treated with SOM alone or in combination with TTX, CYN154806 or Cyclosom (as indicated). B) The bar chart depicts the mean amplitude of the SOM-induced outward currents under the different recording conditions shown in A). The numbers in brackets indicate the numbers of recorded cells. Mean outward current SOM: 33.87 ± 16.78 pA, SOM+TTX: 28.47 ± 11.28 pA, SOM+CYN: 6.73 ± 4.98 pA, SOM+Cylcosom: 13.69 ± 8.56 pA SOM vs. SOM+CYN: *p*<0.001; SOM vs. SOM+Cyclosom: *p*<0.01; SOM+TTX vs. SOM+CYN: *p*<0.01, SOM+TTX vs. SOM+Cyclosom: *p*<0.05; One-way ANOVA with Tukey’s Multiple Comparison Test. C) Representative current recording of a PC receiving voltage ramps at a frequency of 0.1 Hz after sequential addition of SOM and baclofen to the bath. D) The bar chart shows the mean drug-induced outward currents. Numbers in brackets indicate the numbers of recorded cells. Mean outward current SOM: 33.87 ± 16.78 pA, SOM+Bac: 86.96 ± 35.89 pA, Bac: 74.4 ± 23.24 pA, Bac+SOM: 78.2 ± 26.46 pA. SOM vs. Bac: *p*<0.001; SOM vs. SOM+Bac: *p*<0.001, SOM vs. Bac+SOM: *p*<0.001, One-way ANOVA with Tukey’s Multiple Comparison Test. E) Left panel: Voltage command ranging from −120 to −20 mV. Right panel: Representative single voltage ramp-induced current traces (average of five traces) of a recording from a PC before exposure to SOM (con, black trace), after bath application of SOM (SOM, green trace) and after addition of baclofen (SOM+baclofen, red trace). F) Pooled I-V relationships of drug-induced currents show that the drug-induced current reverses between −90 and −80 mV (control: n=19, SOM: n=11; SOM+baclofen: n=8; baclofen: n=5). G) Pooled I-V relationships of PCs recorded in the presence of BaCl2 (black trace) and after application of SOM (green trace). Pre-treatment with BaCl2 blocked the SOM-induced outward current (SOM: n=10, BaCl2: n=11).

Previous studies have reported that the GABA_B_R agonist baclofen also induces an outward current in neurons [33, 34]. Therefore, we compared the magnitude of the SOM-with that of the baclofen-mediated outward current in PCs. To this end, we applied either SOM or baclofen or a sequential application of either agonist (Fig 1C). We observed the following effects: 1) Baclofen induced a significantly larger outward current compared to SOM (Fig 1 C, D), indicating a higher intrinsic efficacy of the drug. 2) SOM failed to induce an additional outward current after pretreatment of slices with baclofen. 3) In contrast, baclofen was still able to induce an additional outward current in PCs that had been pre-exposed to SOM (Fig 1 C, D). These data indicate that baclofen displays a higher intrinsic efficacy compared to SOM and possibly leads to a desensitization of SOM-induced outward currents.

By applying a ramp protocol (ranging from a membrane holding potential of −120 mV to − 20 mV), we then confirmed that the SOM- and baclofen-mediated outward currents were driven by an increased potassium conductance [8, 33–39], as the drug-induced current displayed a reversal potential ranging between −80 mV and −90 mV and was blocked by low concentrations of Barium ions [300 µM] (Fig 1 E-G). These results confirm that baclofen and SOM both increase potassium conductance via GIRK channels in L2/3 PCs.

### 2.2 SOM reduces glutamatergic neurotransmission by depressing presynaptic glutamate release

Having shown that SOM decreases PC excitability by acting on GIRK channels, we next wanted to test whether the neuropeptide would modulate spontaneous synaptic activity in PCs. Therefore, we initially recorded spontaneous excitatory postsynaptic currents (sEPSCs) before and during bath application of SOM in the presence of the GABA_A_R blocker bicuculline. To this end, cells were clamped at a holding potential of −65 mV and bicuculline was added to the bath at least 15 min prior to baseline recording. We found that bath application of SOM had no effect on sEPSC amplitude but caused a significant reduction in sEPSC frequency (Fig 2 A-E), indicating that SOM decreased presynaptic glutamate release in excitatory afferents terminating on PCs.

**Fig 2.**
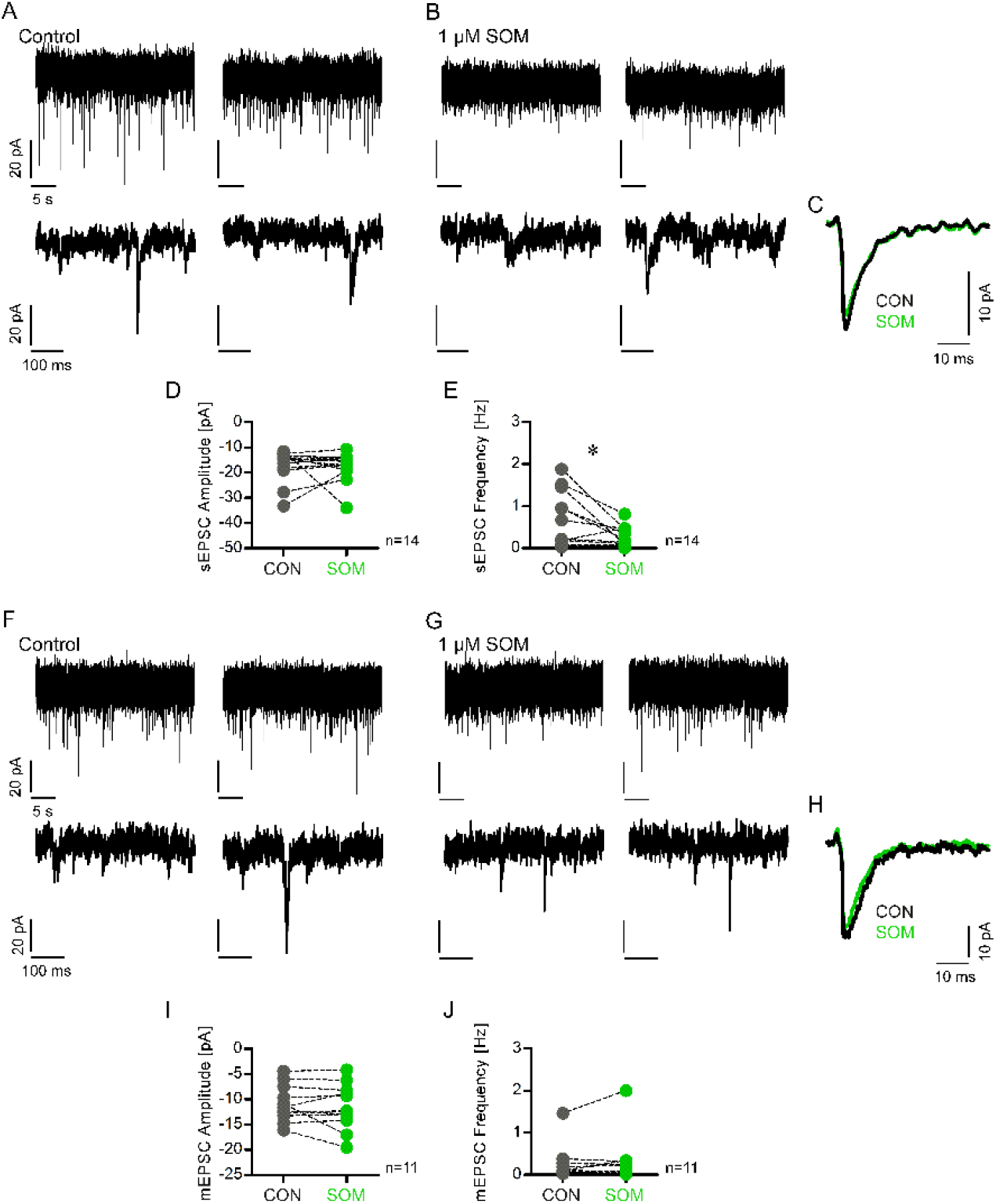
SOM depresses spontaneous excitatory transmission in L2/3 PCs. A) Upper panel: Two representative current traces recorded from a PC under control condition showing spontaneous excitatory postsynaptic currents (sEPSCs) in the presence of bicuculline. Lower panel: Depiction of expanded current traces f the same cell shown in the upper panel. B) Upper panel: Two individual current traces of the same PC as in A) after bath application of 1 µM SOM. Lower panel: Expanded current traces of the corresponding traces in the upper panel. C) Average sEPSC trace of all events detected under control condition (black trace) and after SOM treatment (green trace) in the same neuron. D) The scatter dot plot depicts the amplitude of all sEPSCs recorded before and after SOM bath application. The data were derived from 14 PCs. Mean sEPSC amplitude control: −17.13 ± 6.14 pA, mean sEPSC amplitude SOM: −17.31 ± 5.64 pA, *p*=0.542, Wilcoxon signed rank test. E) The scatter dot plot depicts all sEPSC frequencies before and after addition of SOM. The data were derived from 14 PCs. Mean sEPSC frequency control: 0.59 ± 0.65 Hz, mean sEPSC frequency SOM: 0.24 ± 0.24 Hz, p=0.028, paired t-test. F) Upper panel: Depiction of two representative current traces of miniature EPSC (mEPSC) recordings under control conditions and in the presence of bicuculline and TTX. Lower panel: Expanded current traces of the corresponding traces shown in the upper panel to depict individual mEPSCs. G) Upper panel: Two representative current traces of mEPSC recordings of the same neuron as in F) in the presence of SOM. Lower panel: Expanded current traces of the corresponding traces in the upper panel. H) Average mEPSC trace of all events detected under control condition (black trace) and after addition of SOM (green trace) in the same neuron. I) The scatter dot plot shows all amplitudes of mEPSCs recorded before and after SOM bath application. The data were derived from 11 PCs. Mean mEPSC amplitude control: −10.82 ± 3.64 pA; mean mEPSC amplitude SOM: −11.44 ± 4.61 pA, *p*=0.41, paired t-test. J) The scatter dot plot depicts the mEPSC frequencies before and after addition of SOM. The data were derived from 11 PCs. Mean mEPSC frequency control: 0.25 ± 0.41 Hz, mean mEPSC frequency SOM: 0.31 ± 0.57 Hz, *p*=0.859, Wilcoxon signed rank test.

Next, we probed endogenous SOM release and its effect on spontaneous excitatory neurotransmission by analyzing the sEPSC frequency and amplitude under control conditions and in the presence of the pan-SSTR blocker Cylcosomatostatin or the specific SSTR2 blocker CYN154806. Bath application of either antagonist had no effect on the sEPSC frequency nor amplitude, suggesting that 1) SOM is not endogenously released in acute brain slices or that 2) PC activity in acute brain slices is too little to disclose the effect of endogenous SOM release on spontaneous excitatory transmission (S1 Fig).

In line with above hypothesis that SOM relies on neuronal activity to inhibit glutamatergic synaptic transmission, we found that SOM neither affected the frequency nor the amplitude of miniature EPSCs (mEPSCs) recorded in the presence of TTX (Fig 2 F-J).

We next asked whether SOM had a similar effect on evoked glutamatergic transmission recorded from PCs. Importantly, under our experimental conditions given, L2/3 PCs mainly receive local and corticocortical input as thalamic projections to the cortex are cut during the slice preparation. Corticocortical afferents reach L2/3 PCs predominantly via cortical layer L1 (L1). We therefore focused on SOM-induced effects on evoked responses in PCs following L1 stimulation.

Accordingly, we positioned a monopolar glass electrode in L1 perpendicular to the recorded PCs in cortical layers 2/3 (L2/3, Fig 3A) and recorded evoked excitatory postsynaptic currents (eEPSCs) in the presence of bicuculline at a holding potential of - 65 mV. After a baseline recording of 10 min, we either applied SOM or Cyclosomatostatin to the bath for 10 min and continued the recording. We found that bath application of SOM significantly reduced the amplitude of pharmacologically isolated eEPSCs (Fig 3 C-E) whereas bath application of Cylcosomatostatin had little effect on the eEPSC amplitude (Fig 3 B, D, E). In order to decipher whether this SOM-induced decrease in eEPSC amplitude was due to a postsynaptic depression of glutamatergic currents or not, we synaptically isolated PCs by bath application of TTX and focally applied glutamate to the apical tuft of L2/3 PC dendrites (Fig 3 F, G). To this end, PCs were clamped at a holding potential of −65 mV and glutamate was puffed at an interval of 180 s after having ensured before that the glutamate-driven response had returned to baseline levels at this application interval. Glutamate was applied three times under control conditions (i.e. in the presence of TTX) before the addition of SOM to the bath. Under these conditions, SOM failed to reduce the amplitude of the glutamate-induced response rendering a postsynaptic SOM effect on glutamate receptors highly unlikely.

**Fig 3.**
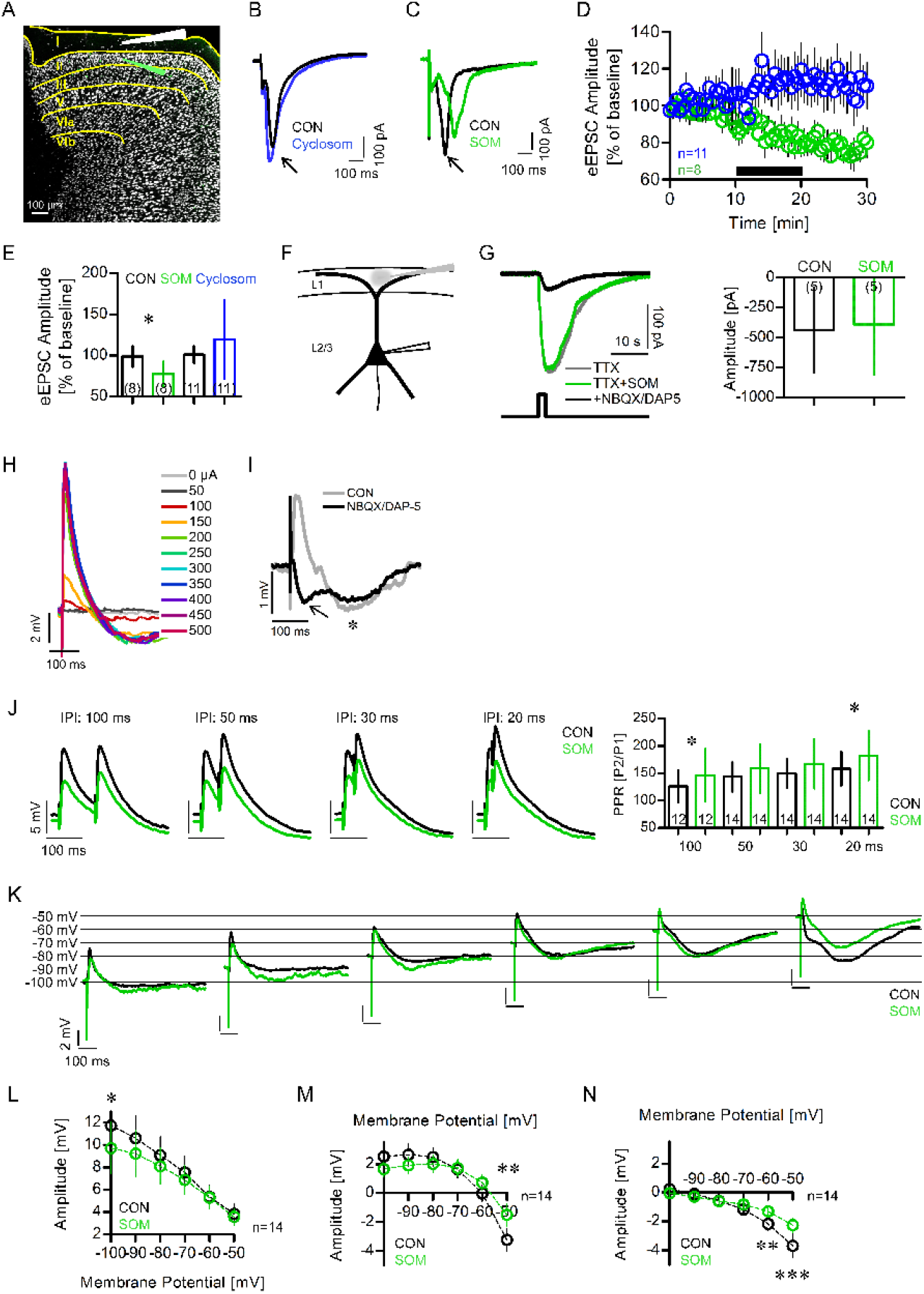
SOM depresses glutamatergic transmission by depressing presynaptic glutamate release. A) Schematized experimental setup. B) Representative single traces (average of three) of evoked excitatory postsynaptic currents (eEPSCs) during baseline recording (black trace) and after addition of Cyclosom (blue trace) to the bath. C) Representative single traces of eEPSCs during baseline recording (black trace) and after addition of SOM (green trace) to the bath. D) The mean eEPSC amplitude plotted as a function of time. E) Bar chart showing the mean normalized baseline just before drug application in comparison to that 10 min after addition of drug. The data were derived from 8 cells (mean normalized eEPSC amplitude con: 98.8 ± 12.52, SOM: 77.6 ± 14.7; *p*= 0.0265, paired t test; mean normalized eEPSC amplitude con: 101.0 ± 10.3, Cyclosom: 119.5 ±47.9 *p=*0.2061, Wilcoxon signed rank test). The numbers in brackets indicate the numbers of recorded cells. F) Schematized experimental setup. G) Left panel: Single representative current traces (average of two) upon focal application of glutamate in the presence of TTX before (grey trace) or after (green trace) bath application of SOM. NBQX/D-AP5 (black trace) was added to show inhibition of glutamate-induced current. Right panel: Bar chart showing the mean glutamate-induced current responses under control condition and after addition of SOM. The data were derived from 5 cells (Mean current amplitude control: −439.5 ± 355.3 pA; SOM: −391.8 ± 414.7 pA, *p*=0.438, Wilcoxon signed rank test). H) Single evoked postsynaptic potentials (ePSPs) of a L2/3 PC after L1 stimulation with different stimulation intensities (average of three). I) Single traces (average of three) of ePSPs recorded in the same PC in the absence (grey trace) or presence (black trace) of 10 µM NBQX and 20 µM D-AP5. J) Left panel: Single ePSP traces (average of three) after paired-pulse stimulation at different interpulse intervals (IPIs) before (black) and after (green) SOM bath application. Right panel: Mean paired-pulse ratio (PPR) under control condition and after SOM bath application. Numbers in bar chart represent numbers of recorded cells (Mean PPR ratio at IPI 100 ms: Control, 126 ± 29.33; SOM, 146 ± 48.54. IPI 50 ms: Control, 143.4 ± 27.27; SOM, 158.7 ± 44.84. IPI 30 ms: Control, 149.6 ± 25.98; SOM, 167.3 ± 45.41. IPI 20 ms: Control, 158.4 ± 31.41; SOM, 182.4 ± 45.21). SOM increases the PPR at an IPI of 100 and 20 ms (IPI 100 ms control vs. SOM, *p*=0.017, Wilcoxon signed rank test; IPI 50 ms control vs. SOM, *p*=0.221, Wilcoxon signed rank test; IPI 30 ms control vs. SOM, *p*=0.055, Wilcoxon signed rank test; IPI 20 ms control vs. SOM, *p*=0.038, paired t test). K) Representative single traces (average of three) of ePSPs in the same L2/3 PC that was held at different membrane potentials before (black trace) and after (green trace) addition of SOM. L) The mean amplitude of the initial fast depolarization under control condition (black) and after addition of SOM (green) plotted as a function of the membrane potential. The data were derived from 14 cells (mean ± SEM; mean ePSP amplitude −100 mV: Control, 11.74 ± 2.25 mV; SOM, 9.71 ± 2.24 mV, p<0.05, Two-way ANOVA with Bonferroni posttest. −90 mV: Control, 10.60 ± 2.04 mV; SOM, 9.22 ± 2.07 mV. −80 mV: Control, 9.08 ± 1.66 mV; SOM, 8.09 ±1.64 mV. −70 mV: Control, 7.54 ± 1.44 mV; SOM, 6.88 ± 1.29 mV. −60 mV: Control, 5.37 ± 1.08 mV; SOM, 5.29 ± 1.00 mV. −50 mV: Control, 3.84 ± 0.90 mV; SOM, 3.57 ± 0.75 mV). M) The mean amplitude of the presumed GABAAR-mediated response under control condition (black) and after addition of SOM (green) plotted as a function of the membrane potential. The data were derived from 14 cells (mean ± SEM; mean ePSP amplitude −100 mV: Control, 2.53 ± 0.74 mV; SOM, 1.70 ± 0.50 mV. −90 mV: Control, 2.66 ± 0.8 mV; SOM, 1.92 ± 0.58 mV. −80 mV: Control, 2.49 ± 0.65 mV; SOM, 2.02 ± 0.60 mV. −70 mV: Control, 1.65 ± 0.60 mV; SOM, 1.74 ± 0.58 mV. −60 mV: Control, −0.03 ± 0.59 mV; SOM, 0.73 ± 0.53 mV. − 50 mV: Control, −3.22 ± 0.77 mV; SOM, −1.49 ± 0.79 mV, *p*<0.001, Two-way ANOVA with Bonferroni posttest). N) The mean amplitude of the presumed GABABR-mediated response under control condition (black) and after addition of SOM (green) plotted as a function of the membrane potential. The data were derived from 14 cells (mean ± SEM; mean ePSP amplitude −100 mV: Control, 0.23 ± 0.10 mV; SOM, −0.05 ± 0.14 mV. −90 mV: Control, −0.10 ± 0.12 mV; SOM, −0.3 ± 0.10 mV. −80 mV: Control, −0.57 ± 0.15 mV; SOM, −0.59 ± 0.12 mV. −70 mV: Control, −1.18 ± 0.24 mV; SOM, −0.83 ± 0.21 mV. −60 mV: Control, −2.17 ± 0.45 mV; SOM, −1.30 ± 0.33 mV; *p*<0.01, Two-way ANOVA with Bonferroni posttest. −50 mV: Control, −3.68 ± 0.84 mV; SOM, −2.26 ± 0.52 mV, *p*<0.001, Two-way ANOVA with Bonferroni posttest).

In order to corroborate a presynaptic mechanism of SOM-induced eEPSC depression, we analyzed the presynaptic facilitation rate as a read-out for presynaptic depression of neurotransmitter release [40]. Here, we analyzed the rate of presynaptic facilitation by recordings of evoked postsynaptic potentials (ePSPs) after electrical stimulation of L1 using paired pulses. Input-output curves were obtained prior each experiment in order to adjust the stimulation intensity to around 80% of the maximum intensity (Fig 3 H). When keeping the PCs at a membrane potential of −60 mV, we found that L1 stimulation evoked an initial, fast depolarizing response that was followed by a biphasic hyperpolarizing response with a faster and a slower onset (Fig 3 I light grey trace). L1 stimulation in the presence of blockers of glutamatergic neurotransmission disclosed that biphasic hyperpolarizing response (Fig 3 I black trace), indicating that L2/3 PCs are under strong inhibitory control via L1. In agreement with this finding, we observed that L1 stimulation in the presence of bicuculline produced prolonged eEPSC responses in L2/3 PCs (Fig 3 B, C, arrow), similar to what has been reported by Shlosberg et al. [41] in L5 PCs after removal of L1 in acute brain slices, indicating that L1 provides strong inhibitory control of L2/3 PCs.

Therefore, given that the evoked postsynaptic response was composed of different synaptic components, we used a paired-pulse stimulation paradigm with different interpulse intervals (IPIs) ranging from 20 ms to 100 ms and analyzed the rate of presynaptic facilitation of ePSPs in the absence or presence of SOM. In agreement with our observation that the SOM-induced depression of eEPSC amplitudes was not due to a depression of the postsynaptic glutamate response itself, we found that SOM increased the paired-pulse ratio (PPR) at an IPI of 20 ms but not at IPIs of 30 and 50 ms (Fig 3 J). In addition, we found that SOM increased the PPR at an IPI of 100 ms.

Having observed that L2/3 PCs are under strong inhibitory control when stimulated in L1, we next recorded ePSPs and simultaneously injected continuous direct current into the recorded neurons to keep them at defined membrane potentials (ranging from −100 to −50 mV, Fig 3K). This analysis allowed us to determine the nature of these ePSP components by plotting the amplitudes of each postsynaptic response component as a function of the respective membrane potential. In doing so, we could show that SOM treatment resulted in a depression of the initial excitatory postsynaptic response (Fig 3 L) at high membrane potentials. In addition, we found that the two distinct hyperpolarizing responses had reversal potentials between −65 and −55 mV and between −100 and −90 mV respectively, indicative of a presumed GABA_A_R- and a presumed GABA_B_R-mediated response (Fig 3 M, N). Bath application of SOM led to a decrease of either inhibitory response at depolarized membrane potentials suggesting that SOM modulates not only glutamatergic but also GABAergic neurotransmission in the aCC.

With regard to the fact that SOM increased the PPR not only at 20 ms IPI but also at 100 ms IPI and that evoked GABAR-mediated postsynaptic responses have a longer latency, it is therefore likely that SOM-induced changes in presynaptic GABA release contribute to the PPR rate at an IPI of 100 ms. (Fig 3 J).

Taken together, we have shown here that SOM depresses spontaneous and evoked excitatory neurotransmission by depressing presynaptic glutamate release. Likewise, our initial data suggest that SOM modulates GABAergic transmission at the presynaptic level. In order to confirm these initial results, our next aim was to pharmacologically distinguish whether SOM affected GABA_A_R- and/or GABA_B_R-mediated synaptic transmission. Therefore, we recorded spontaneous GABA_A_R-mediated inhibitory currents and evoked GABA_A_R- and GABA_B_R-mediated synaptic responses in the absence or presence of SOM.

### 2.3. SOM modulates spontaneous GABAergic synaptic transmission in L2/3 PCs and in SOM-INs and L1-INs

We next investigated whether SOM had any effect on spontaneous GABAergic synaptic transmission onto PCs. Therefore, we pharmacologically isolated spontaneous inhibitory postsynaptic currents (sIPSCs) by addition of NBQX and D-AP5 to the bath. Recordings were performed using a high-chloride internal solution resulting in a chloride equilibrium potential close to 0 mV. PCs were clamped at a holding potential of −65 mV. We analyzed the frequency and amplitude of sIPSCs in the absence or presence of SOM and found that bath application of SOM had no effect on the sIPSC amplitude but significantly increased the sIPSC frequency in PCs (Fig 4 A-E). This unexpected finding suggests that SOM enhances GABAergic inhibition of PCs most likely by activating inhibitory INs that are located presynaptically to PCs. However, similar to glutamatergic miniature currents, SOM had no impact on the mIPSC amplitude nor frequency (Fig 4 F-J) suggesting an activity-dependent effect.

**Fig 4.**
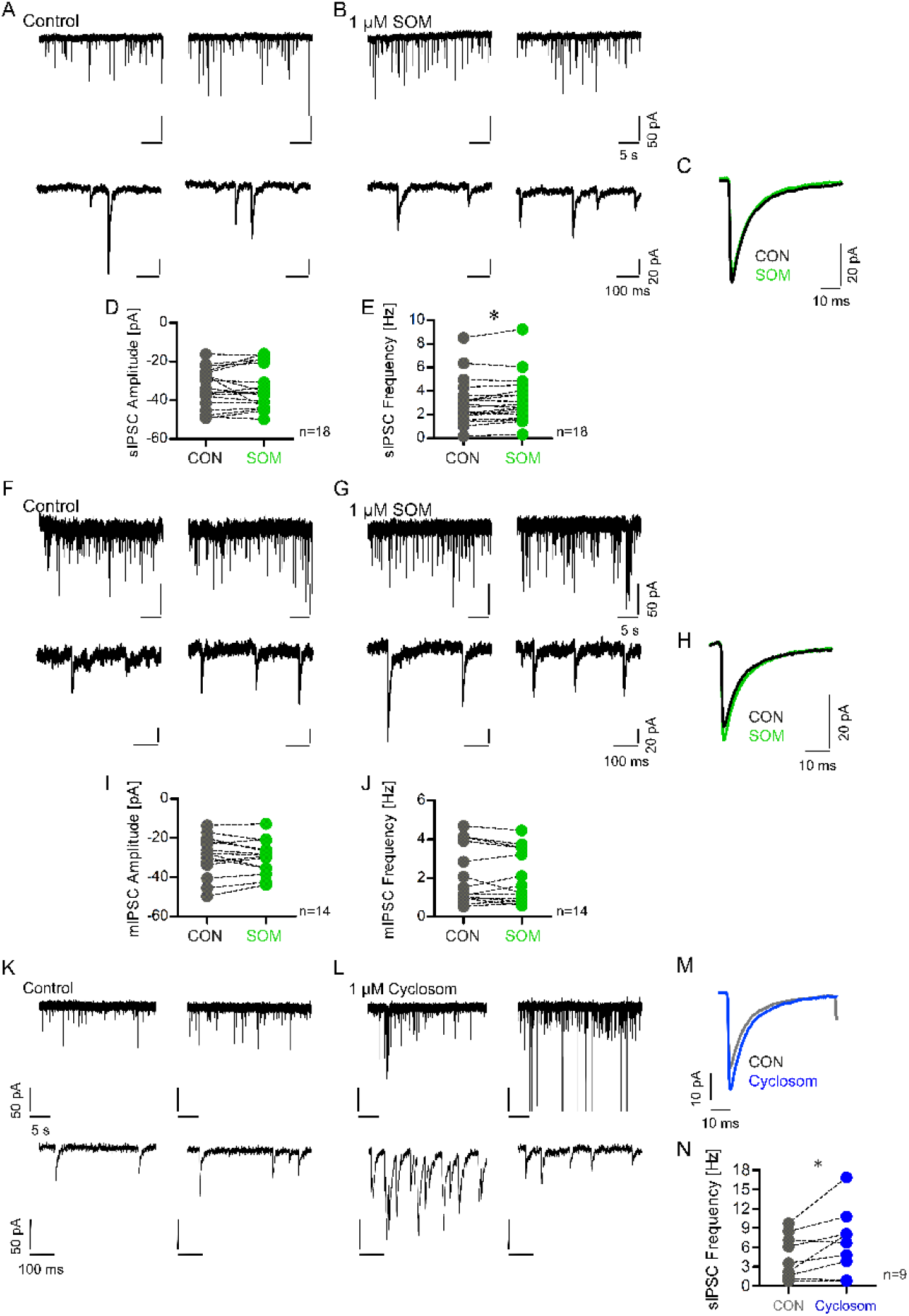
SOM increases the sIPSC frequency in L2/3 PCs. A) Upper panel: Two representative current traces a PC under control conditions showing spontaneous inhibitory postsynaptic currents (sIPSCs). Lower panel: current trace of the corresponding traces in the upper panel to depict individual inhibitory events. B) Upper panel: Two individual current traces of the same PC as in A) after bath application of SOM. Lower panel: Expanded current traces of the corresponding traces in the upper panel. C) Average sIPSC trace of all events detected under control condition (black trace) and after SOM treatment (green trace) in the same neuron. D) The scatter dot plot depicts the mean amplitudes of all sIPSCs recorded before and after SOM bath application. The data were derived from 18 PCs. Mean sIPSC amplitude control: −33.94 ± 10.04 pA, mean sIPSC amplitude SOM: −33.96 ± 11.06 pA, *p*=0.99, paired t-test test. E) The scatter dot plot depicts the mean sIPSC frequencies before and after addition of SOM. The data were derived from 18 PCs. Mean sIPSC frequency control: 3.06 ± 2.00 Hz, mean sIPSC frequency SOM: 3.26 ± 2.04 Hz, *p*=0.029, Wilcoxon signed rank test. F) Upper panel: Depiction of two representative current traces of mIPSC recordings under control condition. Lower panel: Expanded current traces of the corresponding traces shown in the upper panel to show individual miniature events. G) Upper panel: Two representative current traces of mIPSC recordings of the same neuron as in F) in the presence of SOM. Lower panel: Expanded current traces of the corresponding traces in the upper panel. H) Average mIPSC trace of all events detected under control condition (black trace) and after addition of SOM to the bath (green trace) in the same neuron. I) The scatter dot plot shows the mean amplitude of all mIPSCs recorded before and after SOM bath application. The data were derived from 14 PCs. Mean mIPSC amplitude control: −29.28 ± 10.43 pA; mean mIPSC amplitude SOM: −30.02 ± 8.64 pA, *p*=0.52, paired t-test. J) The scatter dot plot depicts the mean mIPSC frequencies of all cells before and after addition of SOM. The data were derived from 14 cells. Mean mIPSC frequency control: 2.12 ± 1.5 Hz, mean mIPSC frequency SOM: 2.04 ± 1.37 Hz, *p*=0.46, paired t-test. K) Upper panel: Depiction of two representative current traces of sIPSC recordings from a L2/3 PC. Lower panel: Expanded current traces of the corresponding traces shown in the upper panel to show individual sIPSCs. L) Upper panel: Two representative current traces of sIPSC recordings of the same neuron as in K) in the presence of Cyclosomatostatin (Cyclosom). Lower panel: Expanded current traces of the corresponding traces in the upper panel. M) Average sIPSC trace of all events detected under control condition (grey trace) and after addition of Cyclosom (blue trace) to the bath in the same neuron. N) The scatter dot plot depicts the mean sIPSC frequencies of all recorded PCs before and after addition of Cyclosom. The data were derived from 9 PCs. Mean sIPSC frequency control: 4.52 ± 3.40 Hz, mean sIPSC frequency Cyclosom: 6.75 ± 5.05 Hz, *p*=0.036, Paired t test.

SOM is thought to be predominantly released by SOM-INs that exert GABAergic inhibition on PCs. Therefore, we next investigated the possible action of endogenously released SOM on sIPSCs recorded in PCs by adding the non-selective SSTR antagonist Cyclosomatostatin or the specific SSTR2 antagonist CYN154806 to the bath. Unexpectedly, we found that either blocker caused a significant increase in the sIPSC frequency in PCs (Fig 4 K-N; S2 Fig), similar to the observed SOM-induced increase in sIPSC frequency. This finding suggests that blockage of endogenous SOM release enhances GABAergic transmission in a subset of GABAergic INs that are located presynaptic to PCs and whose activity is controlled by SOM-INs. In addition, it suggests that the GABAergic INs responsible for the sIPSCs recorded in PCs are differentially affected by SOM.

In order to shed light on this hypothetical cell-type specific action of SOM, we next recorded pharmacologically isolated sIPSCs in SOM-INs and L1-INs before and after addition of SOM to the bath. SOM was added to the bath for 5-7 minutes following a 5 min baseline recording. In agreement with above hypothesis that SOM differentially affects GABAergic INs, we found that SOM significantly increased the sIPSC frequency in L1-INs but decreased the sIPSC frequency in SOM-INs (Fig 5). In addition, SOM had no effect on the sIPSC amplitude in L1-INs but increased the latter in SOM-INs (Fig 5).

**Fig 5.**
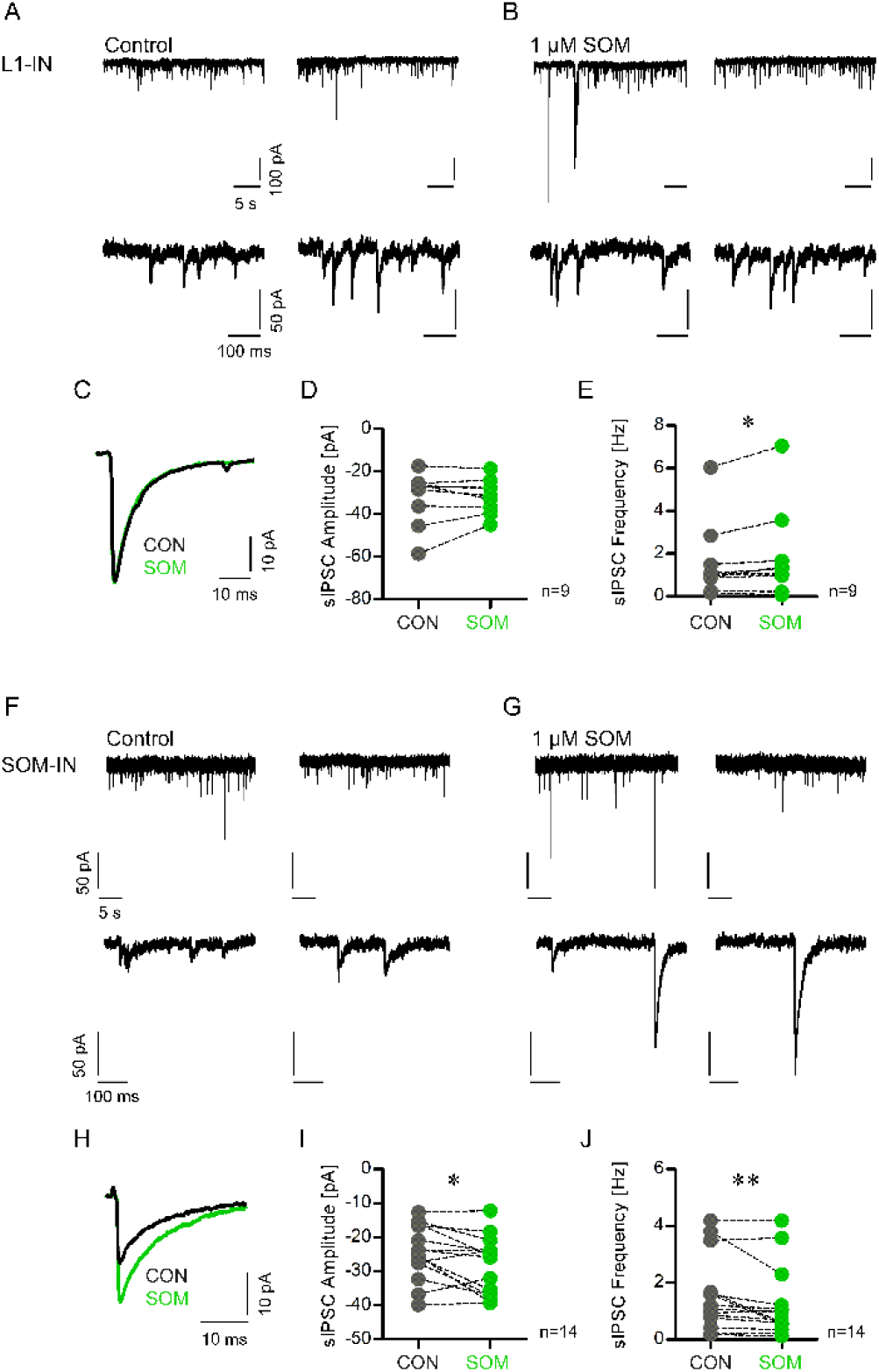
SOM increases the sIPSC frequency in L1-INs but decreases the sIPSC frequency in SOM-INs. A) Upper panel: two representative current traces of a L1-IN under control conditions showing spontaneous inhibitory postsynaptic currents (sIPSCs). Lower panel: current trace of the corresponding current traces of the upper panel to depict individual inhibitory events. B) Upper panel: two individual current traces of the same L1-IN as in A) after bath application of SOM. Lower panel: Expanded current traces of the corresponding traces in the upper panel to show single sIPSCs. C) Average sIPSC trace of all events detected under control condition (black trace) or after SOM treatment (green trace) in the same neuron. D) The scatter dot plot depicts the average amplitudes of all sIPSCs recorded in L1-INs before and after SOM bath application. The data were derived from 9 L1-INs. Mean sIPSC amplitude control: −32.37 ± 12.53 pA, mean sIPSC amplitude SOM: −31.50 ± 8.13 pA, p=0.67, paired t-test test. E) The scatter dot plot depicts the mean sIPSC frequencies before and after addition of SOM. The data were derived from 9 L1-INs. Mean sIPSC frequency control: 1.62 ± 1.83 Hz, mean sIPSC frequency SOM: 1.91 ± 2.17 Hz, *p*=0.039, Wilcoxon signed rank test. F) Upper panel: two representative current recordings of a SOM-IN under control condition. Lower panel: current trace of the corresponding current traces of the upper panel. B) Upper panel: two individual current traces of the same SOM-IN as in F) after bath application of SOM. Lower panel: Expanded current traces of the corresponding traces in the upper panel. C) Average sIPSC trace of all events detected under control condition (black trace) or after SOM treatment (green trace) in the same neuron. D) The scatter dot plot depicts the mean amplitudes of sIPSCs recorded before and after SOM bath application. The data were derived from 14 SOM-INs. Mean sIPSC amplitude control: −24.64 ± 8.01 pA, mean sIPSC amplitude SOM: −28.22 ± 8.47 pA, *p*=0.03, paired t-test test. E) The scatter dot plot depicts the mean sIPSC frequencies before and after addition of SOM. The data were derived from 14 SOM-INs. Mean sIPSC frequency control: 1.57 ± 1.32 Hz, mean sIPSC frequency SOM: 1.24 ± 1.24 Hz, *p*=0.006, Wilcoxon signed rank test.

To strengthen above hypothesis, postsynaptic effects of SOM on L1-INs and SOM-INs were investigated. We limited our analysis on the neuronal circuitry between L2/3 PCs, L1-INs and L2/3 SOM-INs because we legitimately assume that, under our recording conditions, SOM preferentially modulates corticocortical information processing, since all thalamic inputs are interrupted by the slice preparation procedure. The fingerprint electrophysiological properties of these 3 cell types are provided in Table 1 and are depicted in S3 Fig.

**Table 1:** Summary of electrophysiological fingerprint properties of PCs, SOM-INs and L1-INs (Mean ± SD)

| Parameter | PC | SOM-IN | L1-IN |
| --- | --- | --- | --- |
| <u>Passive membrane properties</u> |  |  |  |
|  | n=163 | n=68 | n=51 |
| Capacitance [pF] | 196.0 ± 67.3 | 104.7 ± 35.9 | 80.5 ± 28.9 |
|  | PC >SOM-IN***, PC>L1-IN*** |  |  |
| τ<sub>0</sub> [ms] | 22.2 ± 14.1 | 37.8 ± 23.5 | 19.3 ± 8.9 |
|  | SOM-IN >PC***, SOM-IN>L1-IN*** |  |  |
| τ<sub>1</sub> [ms] | 1.81 ± 2.0 | 4.04 ± 5.1 | 1.72 ± 1.12 |
|  | SOM-IN >PC***, SOM-IN>L1-IN*** |  |  |
| Input Resistance [MΩ] | 150.1 ± 85.9 | 428.0 ± 201.0 | 288.6 ± 131.9 |
|  | SOM-IN>L1-IN**, SOM-IN>PC***, L1-IN>PC*** |  |  |
| <u>Single Spike Properties</u> |  |  |  |
| Rise-to-Fall Ratio | 5.23 ± 1.07 | 2.59 ± 0.47 | 3.12 ± 0.66 |
|  | PC>L1-IN***, PC> SOM-IN***, L1-IN>SOM-IN** |  |  |
| AP Duration [ms] | 1.37 ± 0.37 | 1.06 ± 0.22 | 1.15 ± 0.28 |
|  | PC>L1-IN***, PC>SOM-IN*** |  |  |
| AP Amplitude [mV] | 106.2 ± 9.4 | 83.7 ± 8.3 | 84.0 ± 11.0 |
|  | PC>L1-IN***, PC>SOM-IN*** |  |  |
| AP Threshold [mV] | -41.4 ± 5.5 | -45.0 ± 3.9 | -40.2 ± 4.3 |
|  | SOM-IN>PC***, SOM-IN>L1-IN*** |  |  |

We first tested whether SOM bath application had any effect on the holding current and excitability of SOM-INs and L1-INs in comparison to L2/3 PCs. SOM failed to induce a detectable outward current in SOM-INs, whereas it generated an outward current in L1-INs with an amplitude significantly smaller compared to that observed in PCs (Fig 6 A, B). In agreement with these observations, we found the largest SOM-induced decrease in input resistance (R_N_) in PCs followed by L1-INs. In contrast, SOM had no effect on R_N_ in SOM-INs (data not shown).

**Fig 6.**
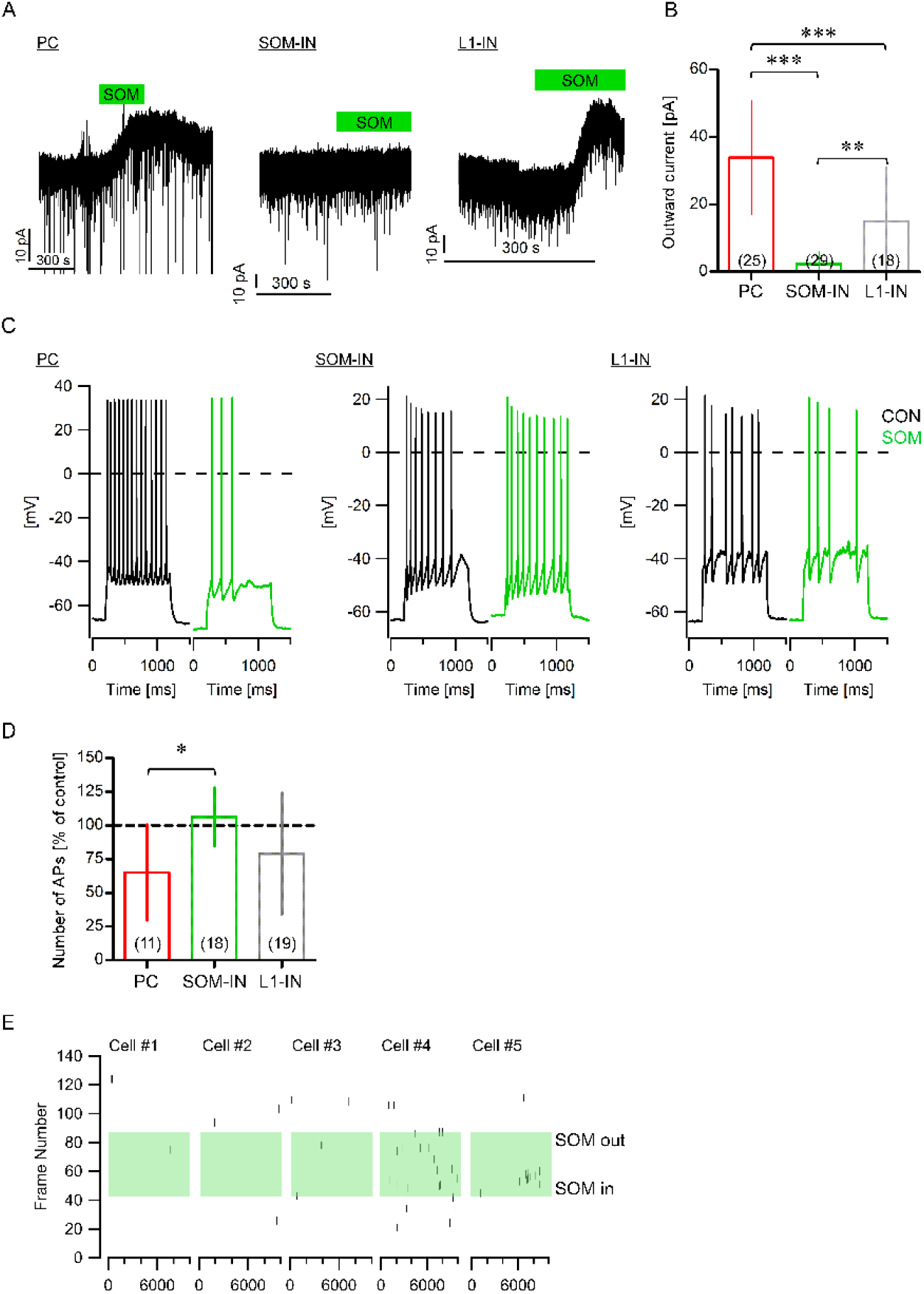
Cell-type specific effect of SOM on outward current in L2/3 PCs, SOM-INs and L1-INs. A) Comparison of SOM-induced outward currents recorded in a L2/3 PC (left panel), in a SOM-IN (middle panel) and in a L1-IN (right panel). B) The bar chart shows the mean amplitude of SOM-induced outward currents. Numbers in brackets indicate numbers of recorded neurons. Amplitude outward current PC: 33.87 ± 16.78 pA, SOM- IN: 2.35 ± 3.53 pA, L1-IN: 14.90 ± 16.37 pA. PC vs. SOM-IN: *p*<0.001, PC vs. L1-IN: *p*<0.001, L1-IN vs. SOM-IN: *p*<0.01, One-way ANOVA with Tukey’s Multiple Comparison Test). C) Single traces of evoked action potentials (APs) upon injection of a supra-threshold depolarizing current step into PCs (left panel), SOM-INs (middle panel) and L1-INs (right panel) before (black trace) and after (green trace) addition of SOM to the bath. D) Summary of the SOM-induced effect on evoked action potential firing in PCs, SOM- INs and L1-INs as bar chart. The numbers in brackets represent the numbers of cells analyzed (mean normalized AP discharge (% of control) PCs: 65.02 ± 35.74, SOM-IN: 106.4 ± 21.85, L1-IN: 79.09 ± 45.09, PC vs. SOM-IN: *p*<0.05, One-way ANOVA with Tukey’s Multiple Comparison Test.). E) Rasterplots showing spontaneous APs discharges in five individual SOM-INs before and after SOM exposure.

In addition, we analyzed whether SOM had any effect on evoked action potential (AP) discharge in the different cell types by injecting a supra-threshold depolarizing current step into the cells before and after bath application of SOM. We found that the SOM- induced action on evoked action potential discharge was significantly different between L2/3 PCs and SOM-INs, supporting above finding that SOM does not negatively affect the excitability of SOM-INs. Moreover, we observed that bath application of SOM increased spontaneous action potential discharge in the majority of SOM-INs, but only in a small fraction of L1-INs or PCs (Fig 6 E).

Taken together, these results suggest that the observed SOM-induced increase in sIPSC frequency in PCs is –at least partially– mediated by an increased activity of presynaptic SOM-INs. Next, we wanted to test whether above finding had any effect on evoked inhibitory synaptic responses in L2/3 PCs.

### 2.4. SOM depresses evoked GABAergic synaptic transmission at the pre- and postsynaptic level in L2/3 PCs

Our next aim was to identify the GABAergic response that was affected by SOM. Therefore, we initially performed an input-output analysis of evoked inhibitory postsynaptic potentials (eIPSPs) recorded from PCs following optogenetic activation of GABAergic fibers in L1 using the VGAT-Chr2-YFP mouse line. PCs were kept at a membrane potential of −60 mV. As mentioned previously, this allowed us to observe a biphasic inhibitory postsynaptic response with an early and a late peak (Fig 7 A, B), indicative of a GABA_A_R- and a GABA_B_R-mediated synaptic component (Fig 7 C). Optogenetic stimulation either by a single pulse or by 10 pulses at 50 Hz revealed that SOM preferentially reduced the late onset GABAergic response in L2/3 PCs (Fig 7 D, E). To distinguish between GABA_A_R- and GABA_B_R-mediated responses, we next isolated the GABA_A_R-driven evoked response by bath application of the GABA_B_R blocker SCH50911. Using the same stimulation paradigm (i.e. either single pulse or 10 pulses at 50 Hz), we found that SOM had no effect on the GABA_A_R-mediated postsynaptic response (Fig 7 F, G). In contrast, using either optogenetic stimulation of evoked GABA_B_R-mediated responses in the presence of bicuculline or using electrical stimulation in the presence of NBQX, D-AP5 and bicuculline, we found that SOM significantly reduced the amplitude of evoked GABA_B_ receptor-driven responses (single pulse or 10 pulses at 50 Hz) in PCs (Fig 7 H-J).

**Fig 7.**
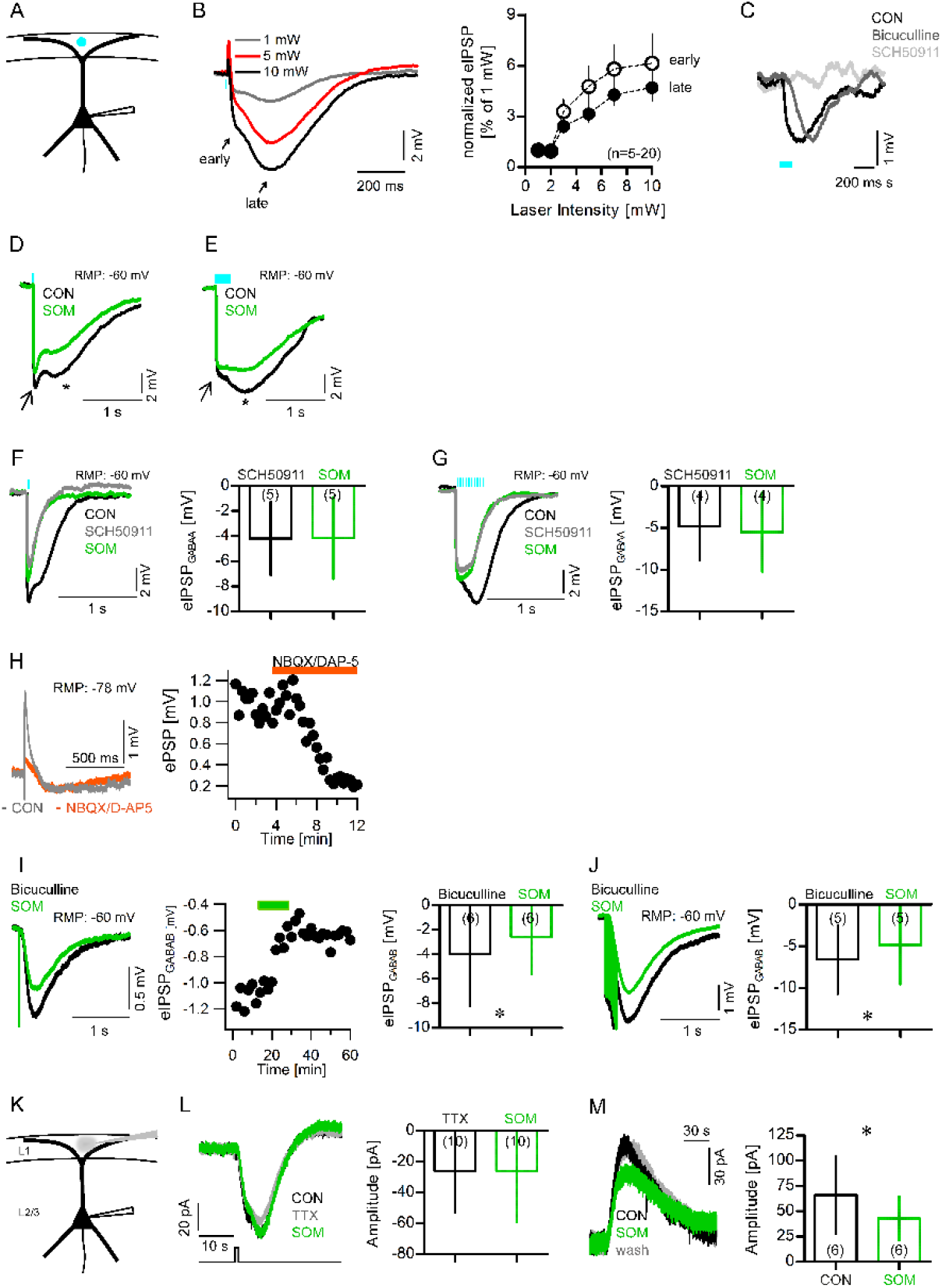
SOM depresses GABAergic synaptic transmission at the pre- and postsynaptic level. A) Schematized recording setup. B) Left panel: Representative single traces (average of three) of optically evoked inhibitory postsynaptic potentials (eIPSPs) in L2/3 PCs at different stimulation intensities. Note the initial fast (early) and the later onset (late) hyperpolarization. Right panel: Input-output curve of the normalized average early and late eIPSP amplitudes at different stimulation intensities. Data were derived from 5-20 L2/3 PCs (mean ± SEM; eIPSP amplitude early: 1 mW, 1.00 ± 0.00 mV (n=20); 2 mW, 0.95 ± 0.33 mV (n=5); 3 mW, 3.29 ± 0.72 mV (n=20); 5 mW, 4.77 ± 1.19 mV (n=18); 7 mW, 5.8 ± 1.45 mV (n=15); 10 mW, 6.15 ± 1.75 mV (n=14). eIPSP amplitude late: 1 mW, 1.00 ± 0.00 mV (n=20); 2 mW, 1.00 ± 0.21 mV (n=5); 3 mW, 2.40 ± 0.39 mV (n=20); 5 mW, 3.15 ± 0.49 mV (n=18); 7 mW, 4.28 ± 0.60 mV (n=15); 10 mW, 4.72 ± 0.81 mV (n=14)). C) Representative single traces (average of three) of optically evoked IPSPs (5 pulses at 50 Hz) under control conditions (black trace), after bath application of bicuculline (dark grey trace) and after bath application of SCH50911 (light grey trace) in the same PC. The cell was kept at a membrane potential of −60 mV. D) Representative optogenetically evoked IPSPs (eIPSPs, average of three) induced by a single light pulse under control (black trace) condition and after bath application of SOM (green trace). Note the initial GABAA receptor- and the late GABAB receptor-mediated response (arrow, resp. asterisk). E) Representative optogenetically evoked IPSPs (eIPSPs, average of three) after 10 light pulses (stimulation frequency 50 Hz) under control (black trace) condition and after bath application of SOM (green trace). Note the initial GABAA receptor- and the late GABAB receptor-mediated response (arrow, resp. asterisk). F) Left panel: Representative single traces (average of three) of evoked GABAAR-mediated postsynaptic potentials before (black) and after bath application of SOM (green) in the presence of SCH50911 following a single laser pulse. Right panel: The bar chart depicts the mean eIPSP amplitudes of the GABAAR-mediated IPSP before (black) and after (green) addition of SOM to the bath. The numbers in brackets indicate the numbers of recorded cells (SCH50911: −4.19 ± 2.93 mV, SOM: −4.15 ± 3.26 mV, *p*=0.921, Paired t test). G) Left panel: Representative single traces (average of three) of evoked GABAAR- mediated postsynaptic potentials before (black) and after bath application of SOM (green) in the presence of SCH50911 following 10 laser pulses at 50 Hz. Right panel: The bar chart depicts the mean eIPSP amplitudes of the GABAAR-mediated IPSPs before (black) and after (green) addition of SOM to the bath. The numbers in brackets indicate the numbers of recorded cells (SCH50911: −4.85 ± 4.07 mV, SOM: −5.54 ± 4.78 mV, *p*=0.19, Paired t test). H) Left panel: Representative ePSP trace (average of three) after electrical stimulation in L1 before (grey) and after addition of NBQX and D-AP5 (orange). The recorded PC had a resting membrane potential (RMP) of −78 mV. Right panel: ePSP amplitude of cell shown in the left panel as a function of time. The orange bar indicates addition of NBQX and D-AP5 to the bath. I) Left panel: Representative single traces (average of three) of the same neuron shown in H). Depiction of evoked GABABR-mediated postsynaptic potentials before (black) and after bath application of SOM (green) in the presence of NBQX, D-AP5 and bicuculline after a single pulse. Middle panel: Amplitudes of eIPSPGABAB of the same neuron are reduced in response to SOM (solid green bar). Right panel: The bar chart depicts the mean eIPSPGABAB amplitudes before (black) and after (green) addition of SOM. The numbers in brackets indicate the numbers of recorded cells (Bicuculline: −3.97 ± 4.24 mV, SOM: −2.57 ± 3.04 mV, *p*=0.0313, Wilcoxon signed rank test). J) Left panel: Representative single traces (average of three) of the same neuron shown in H) and I). Depiction of evoked GABABR-mediated postsynaptic potentials before (black) and after bath application of SOM (green) in the presence of NBQX, D-AP5 and bicuculline following 10 pulses at 50 Hz. Right panel: The bar chart depicts the mean eIPSPGABAB amplitudes before (black) and after (green) addition of SOM to the bath. The number in brackets indicates the number of recorded cells (Bicuculline: − 6.59 ± 4.12 mV, SOM: −4.86 ± 4.73 mV, *p*=0.045, paired t test). K) Schematized experimental setup. L) Left panel: Representative single current traces (average of two) in response to a GABA puff before addition of TTX (CON, black trace), after addition of TTX (grey trace) and after exposure to SOM (green trace). Right panel: Mean GABA puff-induced current responses under TTX (black) condition and after SOM bath application (green) as bar chart. The numbers in brackets indicate the numbers of recorded cells (Mean GABA response TTX: −25.97 ± 26.86 pA, mean GABA response SOM: −26.01 ± 32.85 pA, *p*=0.557, Wilcoxon signed rank test). M) Left panel: Representative single traces (average of two) of baclofen puff-induced currents under control condition in the presence of TTX (CON, black trace), after addition of SOM to the bath (green trace) and after SOM washout (wash, grey). Right panel: Mean baclofen puff-induced current responses under control condition (black) and after SOM bath application (green) as bar chart. The numbers in brackets indicate the numbers of recorded cells (mean baclofen current control: 65.87 ± 38.61 pA, mean baclofen current SOM: 42.74 ± 21.49 pA, *p*=0.0375, Paired t test).

In addition, in order to rule out that SOM acted postsynaptically to potentially modulate GABA_A_ and GABA_B_ receptor-mediated responses, we synaptically isolated PCs by bath perfusion with TTX. GABA was then applied via pressure application to the apical tuft of L2/3 PC dendrites in L1. We chose a puffing interval of 180 s to make sure that the GABA-mediated current had returned to baseline level and that all GABA was washed out from the dendrites prior to the next application. GABA was puffed under control conditions (i.e. in the presence of TTX) and after bath application of SOM. SOM bath application had no effect on the postsynaptic GABA_A_R-mediated response confirming our initial hypothesis that SOM acts presynaptically to modulate GABAergic synaptic transmission (Fig 7 K, L). However, repeated pressure application of the GABA_B_R agonist baclofen at an interval of 240 s triggered an outward current in PCs that became significantly reduced after bath application of SOM suggesting that SOM induced a depression of postsynaptic GABA_B_R-mediated responses (Fig 7 M).

Taken together, our data show that SOM acts pre- and postsynaptically in PCs to modulate GABAergic transmission. Furthermore, we have shown that SOM has a substantially larger effect on PC excitability compared to L1-IN excitability with no detectable effect on SOM-IN excitability. Therefore, we next asked whether above findings would influence neuronal activity within the circuitry between L1-INs and L2/3 PCs and SOM-INs. Thus, we performed recordings of pairs of PCs, pairs of PCs and SOM-INs, pairs of PCs and L1-INs, and pairs of SOM-INs and L1-INs in an effort to first determine the synaptic connections between these cell types and second to understand whether SOM affects correlated activity between these pairs of neurons in an open network, i.e. without any pharmacological intervention.

### 2.5. SOM increases correlated activity between pyramidal cells and decreases correlation between SOM-interneurons and pyramidal cells

Paired recordings between L2/3 PCs and SOM-INs revealed a coupling probability of 17% from PC onto SOM-IN (4 out of 23 pairs recorded, Fig 8 A, B), but synaptic coupling from SOM-INs to PCs could not be detected, possibly due to the fact that SOM-INs innervate the distal dendrites of PCs [17, 18]. However, we found indirect evidence for projections from SOM-INs to PCs by performing paired recordings between PCs. Initially, we determined the coupling probability between pairs of PCs (11% or 7 out of 66 pairs recorded, Fig 8 C, D). Previous studies suggest a similar coupling probability between pairs of L2/3 PCs in the somatosensory [42–45] and visual cortex [46, 47]. Disynaptic inhibition could be recorded in 28.57% (2 out of 7 coupled pairs) of synaptically coupled PCs (Fig 8 E, F) and previous studies suggest that this form is inhibition is mediated by SOM-INs [42]. SOM-INs and L1-INs exhibited a coupling probability of SOM-IN to L1-IN of 12% (2 out of 17, Fig 8 G, H) while the coupling probability of L1-INs to SOM-INs was only 6% (1 out of 17). The coupling probability of L1-INs to PCs was 33% (3 out of 9 pairs) and PCs exhibit a coupling ratio of 22% (2 out of 9 pairs, Fig 8 I, J) with L1-INs. It should be noted here that we recorded from a mixed population of L1-INs and we made no distinction between subtypes of L1-INs. This is reflected in the connection probability between L1-INs to PCs: while the coupling probability of a subtype of L1-INs, namely neurogliaform cells, to PCs ranges from 7% (rat neocortical slices, [48] to 72% (mouse barrel cortex, [49] that of another subtype of L1-INs, namely canopy cells, amounts to 18% [49], i.e. the coupling probability of L1-INs to PCs presented here, is a close average of these coupling probabilities. Taken together, we found the strongest synaptic connections between PCs and L1-INs followed by PCs and SOM-INs (Fig 8 K).

**Fig 8.**
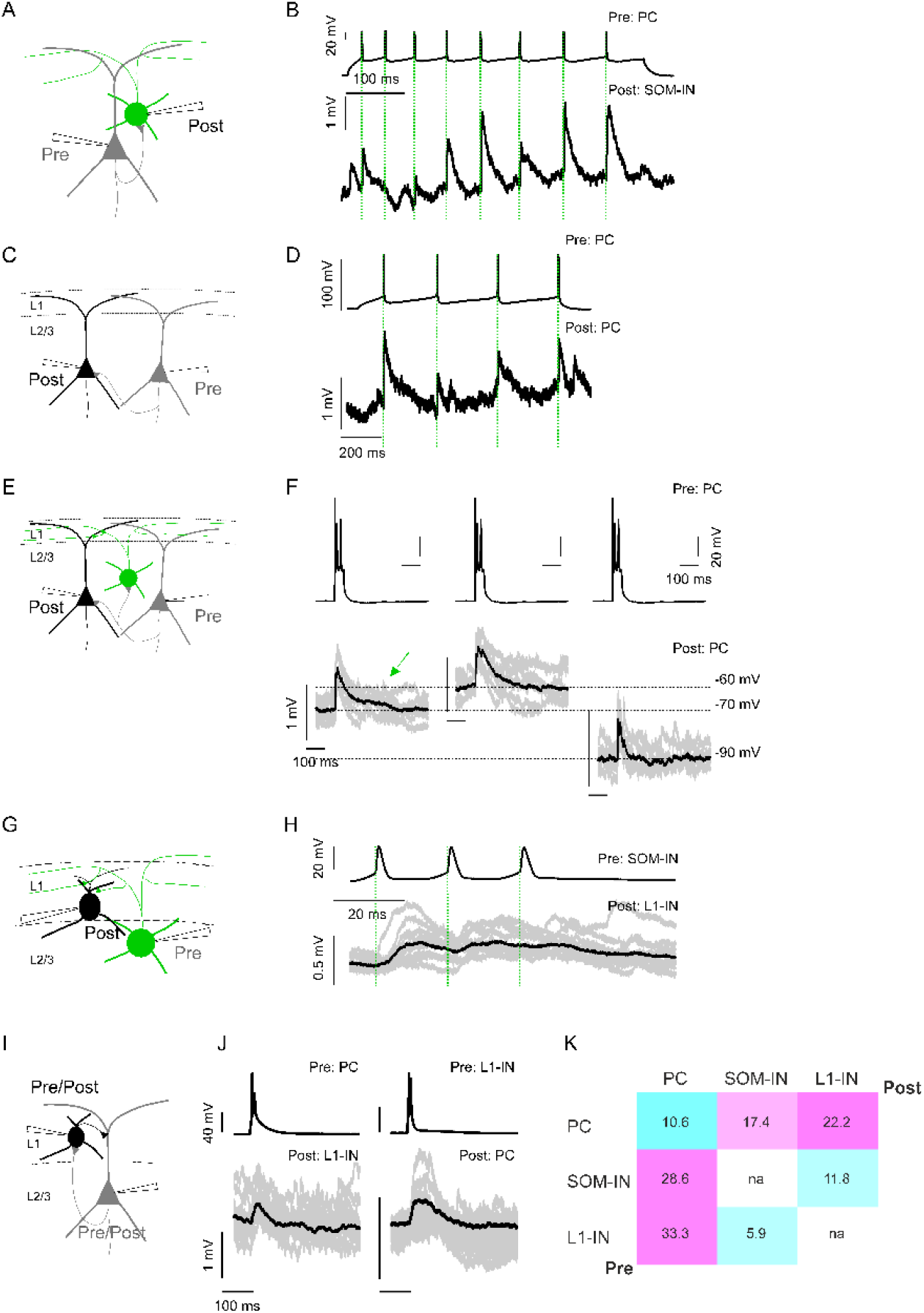
Synaptic connectivity between L2/3 PCs, L2/3 SOM-INs and L1-INs of the aCC. A) Schematized recording setup. B) Presynaptic action potential discharge in a PC evoked unitary EPSPs (uEPSPs) in the postsynaptic SOM-IN. C) Recordings from a pair consisting of PCs. D) Action potentials in the presynaptic PC evoked uEPSPs in the postsynaptic PC. E) Paired recording similar to C) with a putative SOM-IN added to the circuitry. F) Action potential discharge in a presynaptic PC evoked a response in the postsynaptic PC. The postsynaptic cell was held at different membrane potentials to analyze different components of the postsynaptic response. The average postsynaptic trace (black) of the PC held at −70 mV indicates a fast postsynaptic response that is followed by a delayed depolarizing response with a longer latency (indicated by the green arrow). The delayed postsynaptic depolarizing response disappeared upon changing of the membrane potential of the postsynaptic PC to −60 mV. G) Recording situation. H) Action potentials in a presynaptic SOM-IN evoked postsynaptic potentials in the L1-IN. Individual responses are depicted in light grey; the mean postsynaptic response is shown in black. I) Paired recordings between L1-INs and L2/3 PCs. J) Bidirectional coupling between PC and L1-INs. Left panel: Presynaptic action potentials in the PC evoked postsynaptic potentials in the L1-INs. Right panel: Presynaptic action potentials in the L1-IN evoked postsynaptic potentials in the PC. K) Heatmap showing the overall connectivity between L2/3 PCs, L2/3 SOM-INs and L1-INs of the aCC.

Having determined the synaptic connections between PCs, SOM-INs and L1-INs, we next asked ourselves whether SOM had any impact on the correlated neuronal activity between above cell pairs. Therefore, we recorded voltage traces of pairs of these cell types in the presence or absence of SOM and analyzed their correlated membrane potential changes. After a baseline recording of at least 5 minutes, SOM was added to the bath for 5-7 minutes and the recording continued for at least 15 minutes. During intact network activity (i.e. no block of glutamatergic or GABAergic transmission), we found that SOM had no effect on the amplitude, frequency, duration or decay-time of spontaneous synaptic potentials (sPSPs) in all 3 cell types examined (S4 Fig), however, we found SOM-induced changes in correlated membrane potential fluctuations between pairs of PCs and between pairs of SOM-INs and PCs and L1-INs and PCs. In agreement with above finding that pairs of PCs only show a minor synaptic coupling ratio, we failed to observe correlated neuronal activity in neighboring PCs. Application of SOM however, resulted in a significant transient increase in correlated activity between these pairs of cells (Fig 9 A-D).

**Fig 9.**
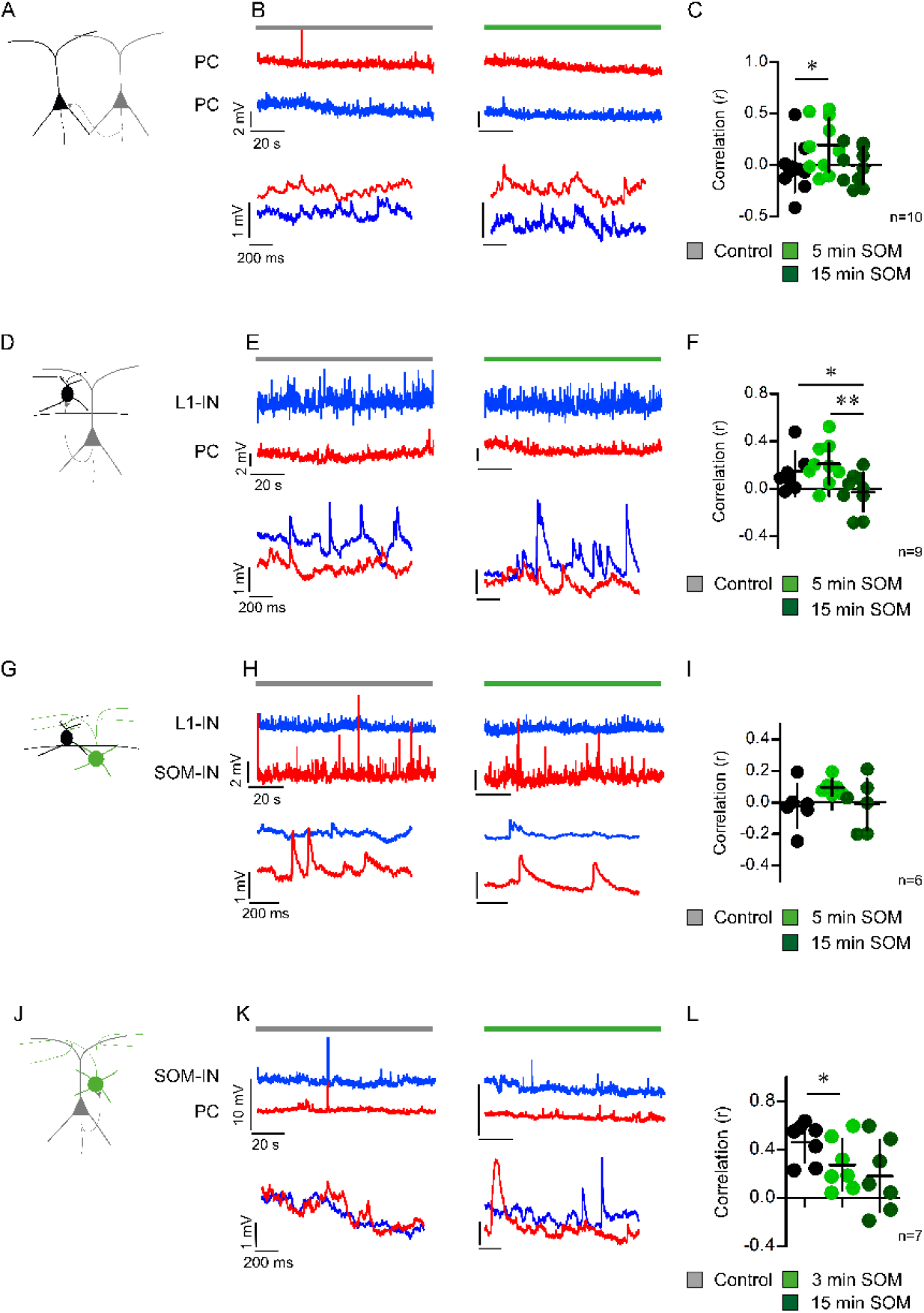
SOM increases correlated activity between pyramidal cells and decreases correlation between SOM-INs and PCs. A) Scheme for recording pairs of L2/3 PCs. B) Upper panel: voltage traces of dual recordings of 2 PCs (red and blue) before (solid grey bar) and after bath application of SOM (solid green bar). Lower panel: voltage traces during baseline recording and around 5 min after addition of SOM to the bath. C) The scatter plot shows the mean Pearson’s linear correlation coefficient under control (grey) condition, and 5 (light green) and 15 min (dark green) after onset of SOM bath application. The data were derived from 10 paired recordings (Mean *r* control: −0.03 ± 0.24, mean *r* 5 min SOM: 0.20 ± 0.26, mean *r* 15 min SOM −0.00 ± 0.18. Control vs. 5 min SOM: p=0.041, paired t-test. Control vs. 15 min SOM: *p*=0.842, paired t test, 5 min SOM vs. 15 min SOM p=0.099, paired t test). D) Recording pairs of L2/3 PC and L1-IN. E) Upper panel: voltage traces of a L1-IN (blue trace) and a PC (red trace) before (solid grey bar) and after bath application of SOM (solid green bar). Lower panel: voltage traces during baseline recording and around 5 min after addition of SOM to the bath. F) The scatter plot shows the mean Pearson’s linear correlation coefficient under control (grey) condition, and 5 (light green) and 15 min (dark green) after onset of SOM bath application. The data were derived from 9 paired recordings (Mean *r* control: 0.15 ± 0.17, mean *r* 5 min SOM: 0.21 ± 0.17, mean *r* 15 min SOM −0.03 ± 0.17. Control vs. 5 min SOM: *p*=0.337, paired t-test. Control vs. 15 min SOM: *p*=0.0157, paired t test. 5 min SOM vs. 15 min SOM: *p*=0.008, Paired t-test). G) Recording pairs of SOM-IN and L1-IN. H) Upper panel: voltage traces of a L1-IN (blue trace) and a SOM-IN (red trace) before (solid grey bar) and after bath application of SOM (solid green bar). Lower panel: voltage traces during baseline recording and around 5 min after addition of SOM to the bath. I) The scatter plot shows the mean Pearson’s linear correlation coefficient under control (grey) condition, and 5 (light green) and 15 min (dark green) after onset of SOM bath application. The data were derived from 6 paired recordings (Mean *r* control: −0.02 ± 0.14, mean *r* 5 min SOM: 0.10 ± 0.05, mean *r* 15 min SOM −0.01 ± 0.16. Control vs. 5 min SOM: *p*=0.170, paired t-test. Control vs. 15 min SOM: *p*=0.90, paired t test. 5 min SOM vs. 15 min SOM: *p*=0.226, Paired t-test). J) Recording pairs of SOM-IN and PC. K) Upper panel: Voltage traces of a SOM-IN (blue trace) and a PC (red trace) before (solid grey bar) and after bath application of SOM (solid green bar). Right panel: voltage traces during baseline recording and around 5 min after addition of SOM to the bath. L) The scatter plot shows the mean Pearson’s linear correlation coefficient under control (grey) condition, and 5 (light green) and 15 min (dark green) after onset of SOM bath application. The data were derived from 7 paired recordings (Mean *r* control: 0.46 ± 0.17, mean *r* 5 min SOM: 0.27 ± 0.21, mean *r* 15 min SOM 0.18 ± 0.29. Control vs. 5 min SOM: *p*=0.032, paired t-test. Control vs. 15 min SOM: *p*=0.062, paired t test. 5 min SOM vs. 15 min SOM: *p*=0.265, Paired t-test).

PC and L1-IN pairs on the other hand showed a moderate degree of correlated activity under control conditions and SOM initially tended to increase correlated activity between these cells but this correlation decreased 15 min after onset of SOM bath application (Fig 9 D-F). In agreement with the fact that different IN subtypes are driven by discrete ensembles of PCs, we could not detect correlated activity between pairs of SOM-INs and L1-INs under control conditions [50]. However, bath application of SOM tended to transiently increase the correlated activity between these 2 cell types (Fig 9 G-I). Lastly, we could show that SOM-INs displayed the highest degree of correlated neuronal activity with PCs and that application of SOM caused a significant reduction of that correlated activity (Fig 9 J-L).

## 3. Discussion

We have shown here that SOM, by acting pre- and postsynaptically, modulates the neuronal circuitry between L1-INs, L2/3 PCs and L2/3 SOM-INs in the aCC. This study revealed a differential effect of SOM on these 3 different cell types and, we could show that SOM failed to reduce SOM-IN intrinsic excitability despite reducing that of PCs and L1-INs. Accordingly, we detected increased sIPSC frequencies in PCs and L1-INs while at the same time, the frequency of inhibitory input onto SOM-INs was reduced in response to SOM treatment. Importantly, we showed that continuous exposure to SOM led to a decrease of GABA_B_R-mediated transmission onto PCs and the data presented here suggest that this decrease is the consequence of a combined pre- and postsynaptic effect on resp. in PCs themselves, due to 1) SOM-induced modulation of presynaptic GABA release and 2) GIRK channel inactivation or desensitization in PCs in response to continuous SOM exposure. Desensitization of SSTRs in response to continuous agonist exposure cannot be entirely ruled out, however, previous work by us points to a desensitization of GIRK channels rather than SSTRs upon ligand binding [8]. Importantly, a similar loss of postsynaptic GABA_B_ receptor-mediated responses was reported in a GIRK knockout mice [37], driving the hypothesis that the combined activation of SSTR and GABA_B_R reduces postsynaptic GIRK channel activity [51].

### Continuous SOM exposure reduces GABA_B_R-mediated signaling and modulates the synaptic strength between INs and PCs

SOM and baclofen both activate GIRK channels in the postsynaptic cell. We found here that the amplitude of the SOM-induced outward current was larger in PCs compared to L1-INs and not detectable in SOM-INs. Cell-type specific baclofen effects were also reported on different hippocampal IN types and in agreement with our findings, baclofen did not induce an appreciable outward current in SOM-INs [52, 53].

The SOM-induced actions on L2/3 PCs seem to be primarily mediated by activation of SSTR2, consistent with previous work by us and others [6–8]. However, detailed analysis of SSTR expression on GABAergic INs revealed prominent expression of SSTR1, SSTR3, SSTR4 and SSTR5 on PV-, SOM- and VIP-INs. Therefore, activation of other SSTR subtypes cannot be ruled out [12]. Given that L1-INs are reported to be the primary mediators of GABA_B_R-mediated inhibition in PCs [54–56], SOM-induced GIRK channel inactivation in PCs results in a smaller L1-IN-mediated inhibition of PCs, i.e. SOM weakens synaptic transmission between L1-INs and PCs. SOM-INs, on the other hand, inhibit PCs to a large extent, but not exclusively, via activation of GABA_A_Rs [43, 54, 57, 58] and we have shown here that SOM specifically weakens GABA_B_R-mediated transmission without having a significant impact on GABA_A_R-mediated transmission, i.e. it is likely that SOM does not affect the synaptic strength between SOM-INs and PCs. It is therefore suggested that SOM-INs, by activity-dependent release of the peptide, gain control over L1-INs and become the predominant regulator of excitatory information flow between PCs. This hypothesis is supported by the finding that optogenetic silencing of cortical SOM-INs enhances synaptic transmission between L2 PCs [43] and increases their action potential firing rate [21].

On a network level, this could potentially mean that lateral inhibition and/or feed-back inhibitory motifs provided by SOM-INs gain importance over feed-forward inhibitory motifs provided by L1-INs upon endogenous release of SOM. Such a scenario would encompass that corticocortical afferents would preferentially activate L1-INs and that SOM-INs only become recruited upon higher activation. Indeed, input-output analyses in PCs, SOM-Ins and L1-INs after electrical L1 stimulation suggest, that L1-INs become recruited well before SOM-INs and PCs (Fig S5). In agreement with such a hypothesis, previous studies using optogenetic stimulation [54] and virus tracing techniques [59], showed that L1-INs received strong corticocortical input.

### SOM modulates dendro-somatic integration in L2/3 PCs

In summary, our findings suggest that, in an open network i.e. with intact GABAergic and glutamatergic transmission, the overall impact of SOM on synaptic excitability of PCs is subtle because increased SOM-induced potassium conductance and decreased GABA_B_R-mediated inhibition in the postsynaptic PC are balanced. In support of this hypothesis, we find that SOM does not alter the input-output ratio of overall evoked postsynaptic responses (data not shown). Likewise, we did not observe a SOM-mediated decrease in ePSP amplitudes in SOM-INs or L1-INs after L1 stimulation (data not shown). It is rather suggested that SOM, by the mechanisms described above, modulates the dendritic integration in apical dendrites of L2/3 PCs. In support of this hypothesis, it could be shown that activation of GABA_B_Rs in apical dendrites of L5 PCs influences the dendro-somatic synergy between feedback and feedforward inputs [55]. We provide evidence here that this influences correlated neuronal activity between pairs of PCs but also between PCs and LI-INs and between L1-INs and SOM-INs while at the same time, decreasing correlated activity between PCs and SOM-INs. *In vivo* recordings of L2/3 PCs and SOM-INs of the barrel cortex during quiet wakefulness showed that membrane potential fluctuations of neighboring PCs are correlated whereas those of PCs and SOM-INs are anti-correlated [21]. In contrast to that, we observed correlated neuronal activity between PCs and SOM-INs under *in vitro* conditions. Correlated membrane potential fluctuations between L2/3 PCs and SOM-INs were also reported by Neske et al [60] in *in vitro* recordings of barrel cortex neurons suggesting that synaptic inputs that are no longer present in *ex vivo* recordings are necessary for anticorrelated activity between SOM-INs and PCs. Previous studies have suggested that SOM-INs play a role in modulating top-down inputs into the cortex [23, 61–64] and behavioral experiments in mice suggest that either bilateral injection of SOM into the visual cortex and/or optogenetic activation of GABAergic long-range efferents from the (posterior) cingulate to the visual cortex enhance visual discrimination and promote center-surround suppression [23, 28]. Likewise, SOM-INs promote sustained spectral surround suppression in excitatory neurons of the auditory cortex [65]. It is therefore suggested that SOM-INs, by endogenous release of the peptide, promote lateral inhibition of PCs. This hypothesis is supported by the fact, that SOM-INs contribute to L1-IN inhibition and recordings from NDNF1 cells suggest, that SOM-INs are important inhibitors of NDNF1-IN activity [54]. We propose here that there exists a disinhibitory SOM-IN-to-L1-IN circuit that possibly gains importance upon endogenous release of SOM.

Ultimately, pre- and postsynaptic effects of SOM on other GABAergic IN types must be incorporated to fully understand how SOM influences the cortical microcircuit and corticocortical information processing as this study did not analyze presynaptic effects of SOM on GABA_B_R signaling in other cortical INs [66]. Given that SOM-INs are reported to inhibit all other IN types [13, 14, 47], differentially modulated pre- and postsynaptic GABA_B_R activation represents an interesting possibility, whereby SOM-INs could regulate GABA release and the strength of synaptic inhibition and inhibitory circuit motifs.

## 4. Materials and Methods

### 4.1. Animals

Experiments were performed on mice from three different lines: 1) a transgenic mouse line (FVB-Tg(GadGFP)45704Swn/J) in which in a subset of SOM-INs expresses the enhanced green fluorescent protein (eGFP) [67], 2) a transgenic mouse line, where all GABAergic INs express channelrhodopsin-2 and YFP (VGAT-ChR2-YFP, [68], and 3) a cross-breed of line 1) and 2). Animals were purchased from Jackson Laboratories (ME, USA) and were bred in the institute’s animal facility. All experiments were approved by the authors’ institutional committees on animal care and were performed according to the German Animal Protection Law, conforming to international guidelines on the ethical use of animals.

### 4.2. Slice preparation

The median age of animals used in this study was postnatal day 27. Animals of either sex were used in the study. The preparation of coronal slices (thickness: 300 µm) was performed as previously described [8, 69]. After the cutting procedure, the slices were collected and submerged in artificial cerebro-spinal fluid (ASCF) containing (in mM): NaCl (125), KCl (3), NaH_2_PO_4_ (1.25), NaHCO_3_ (25), CaCl_2_ (2), MgCl_2_ (2) and D-Glucose (25 mM). The ACSF was continuously perfused with 95% O_2_ / 5% CO_2_ to maintain a pH of 7.4. The osmolarity of the ACSF ranged between 310 to 323 mOsmol. Before recording, the slices were left to recover from the cutting procedure for 30 min at 28°C.

### 4.3. Whole-cell patch-clamp recordings

A single slice was transferred to a recording chamber mounted on the stage of an upright microscope (BX-RFA-1-5, Olympus, Japan). The ACSF temperature was maintained at 27-28°C with the help of a temperature controller (TC-324B, Warner Instrument Corp., Connecticut, USA). All cells were visualized with a Dodt contrast tube (DGC, Scientifica, UK) that was attached to the microscope. SOM-INs were additionally visualized by eGFP expression with the help of a fluorescence lamp (pE-300, CooLED, UK) and epifluorescence optics for green fluorescence (Chroma Technology, USA). Images were taken and displayed using a software-operated microscope camera (Evolve 512 Delta, Teledyne Photometrics, USA). Electrodes were fabricated from borosilicate glass capillaries (OD: 1.5 mm, ID: 0.86 mm, Hugo Sachs Elektronik-Harvard Apparatus, March-Hugstetten, Germany) and were filled with a solution containing (in mM): K-gluconate (135), KCl (4), NaCl (2), EGTA (0.2), HEPES (10), Mg-ATP (4), Na-GTP (0.5), and phosphocreatine (10) or with a solution containing (in mM): KCl (139), NaCl (2), EGTA (0.2), HEPES (10), Mg-ATP (4), Na-GTP (0.5), and phosphocreatine (10). All solutions had an osmolarity of 290-300 mOsm and a pH of 7.3. A silver / silver chloride – pellet served as reference electrode. Somatic whole-cell recordings were made in current- or voltage-clamp mode using an ELC 03XS amplifier (npi electronics, Tamm, Germany). After rupture of the membrane, the electrode capacitance and series resistance were compensated as previously described [70]. The resting membrane potential of all cells ranged between −58 mV and −85 mV.

### 4.4. Electrical or optical stimulation

Stimulation was performed by either placing a monopolar glass electrode into cortical layer 1 or by applying a short focal laser pulse (wavelength: 473 nm, light pulse diameter: 5 µm) onto cortical layer 1 perpendicular to each recorded cell. Stimulation current intensities ranged between 10-500 µA and the pulse duration was 100 µs. Laser intensities ranged between 1-15 mW and lasted for 1-3 ms. Evoked responses were recorded after a single stimulus, after a paired stimulus (2 pulses at 100, 50, 30 and 20 ms interpulse interval (IPI)) or after 10 stimuli (10 pulses at 20 ms IPI). All stimulation protocols were applied at an interval of 15-30 s.

### 4.5. Drug application

Blockers of glutamatergic neurotransmission were: NBQX [10 µM] and D-AP5 [20 µM]. GABA_A_R-mediated transmission was blocked by addition of 10 µM bicuculline. GABA_B_R-mediated signaling was inhibited by SCH50911 [10 µM]. Action potentials were blocked by tetrodotoxin (TTX) at a concentration of 0.5 µM. Somatostatin (SOM), the pan-SSTR antagonist Cyclosomatostatin and the selective SSTR2 antagonist CYN15048 were added at a concentration of 1 µM. All drugs were either applied by addition to the bath or by local pressure application.

#### Bath perfusion

All drugs were either obtained from Tocris Bioscience (Bristol, UK) or from Bio-Trend GmbH (Cologne, Germany). (Antagonists of GABAergic (bicuculline, SCH50911), glutamatergic transmission (D-AP5, NBQX) or SSTR antagonists (Cyclosomatostatin, CYN154806) were added to the bath at least 10 min prior to recording and were left inside the bath until the end of the recording session. In some cases, antagonists were washed out to test reversibility of the effect. SOM and/or baclofen were applied for 5-10 minutes at a concentration of 1 µM (SOM) or 30 µM (baclofen). The volume of the perfusion system was about 9 ml and the perfusion rate was 3 mL/min.

#### Pressure application

GABA [10 mM], glutamate [10 mM] and baclofen [300 µM] were applied via focal pressure application. To this end, a glass capillary was positioned in cortical layer 1 perpendicular to the recorded cell and the pressure was adjusted to 300-500 mbar. The pressure pulse lasted for 1 s and the application interval ranged between 180 s – 240 s.

### 4.6. Data acquisition and analysis

Voltage and current signals were amplified, filtered at 20 kHz (current-clamp recordings) or at 3-5 kHz (voltage-clamp recordings) and digitized at sampling rates of 10-50 kHz. Data acquisition and generation of command pulses was accomplished by an analogue-digital converter (CED 1401 Power 3, Cambridge Electronic Design, Cambridge, UK) in conjunction with the Signal data acquisition software (Version 6, Cambridge Electronic Design). Data analysis was performed using IGOR Pro 9 (WaveMetrics, Lake Oswego, USA) together with the NeuroMatic IGOR plugin (www.neuromatic.thinkrandom.com). Intrinsic electrophysiological properties of recorded neurons were assessed as previously described [69]. SOM- and/or baclofen-induced differences in the holding current were analyzed at a membrane potential of −60 mV with either a K-Gluconate-based or a K-Chloride-based intracellular solution. Spontaneous postsynaptic currents were analyzed at a holding potential of −65 mV and were automatically detected using the algorithm provided by the NeuroMatic plugin (Version 3.0, [71]). In voltage-clamp recordings, the detection threshold was set to −10 pA, in current-clamp recordings, the detection threshold was set to 0.3-0.4 mV. We analyzed the frequency, amplitude, decay-time and rise-time of all synaptic events under control condition (5 min) and after drug application (5 min). Evoked synaptic responses and agonist-induced responses (GABA, glutamate, SOM and baclofen) were assessed by determining the peak voltage or current and by subtracting the baseline voltage or current from this peak value. Synaptic coupling between pairs of neurons was tested by injecting at least 20 supra-threshold depolarizing current steps into the presynaptic neuron and analyzing the corresponding voltage changes in the postsynaptic neuron. Synaptic coupling was defined successful if the average voltage trace of the passive neuron showed a postsynaptic response within a (peak-to-peak) latency of <7 ms. Cross-correlation of neuronal activity between pairs of neurons was quantified in IGOR Pro. To this end, 80-100 s long voltage traces obtained from pairs of neurons under control condition and 5 or 15 min after bath application of 1 µM SOM were analyzed for correlation assessing the Pearson linear correlation coefficient *r* according to Nazemi and Jamali [72]. Statistical analyses were performed by initially testing for a normal distribution of data points using the D’Agostino and Pearson Omnibus Normality test. In case of a normal distribution of data points, paired or unpaired two-tailed students’ *t*-test were performed for comparisons between two different conditions. In case of a non-normal distribution of data points, Mann-Whitney tests or Wilcoxon signed rank tests were performed. Comparisons between multiple conditions were performed by one-way ANOVA or repeated measures ANOVA with Tukey’s Multiple Comparison tests when data points were normally distributed. In case of a non-normal distribution of data points Kruskal-Wallis or Friedman test with Dunn’s Multiple Comparison tests were applied. Significance levels were *p*<0.05 (*), *p*<0.01 (**) and *p*<0.001 (***). Data are presented as mean ± standard deviation, unless otherwise indicated. Graphs were prepared in IgorPro, in GraphPad Prism (La Jolla, CA, USA) or in RStudio (version 1.3.1093, https://www.r-project.org/).

## Acknowledgements

We would like to thank Gabi Horn for excellent technical assistance. This work was funded by 2 grants from the Friedrich-Baur Stiftung (04/16, 03/21).

## Data Availability

All datasets from all figures are provided as Source Data Files.

## Supplemental data

**S1 Fig.**
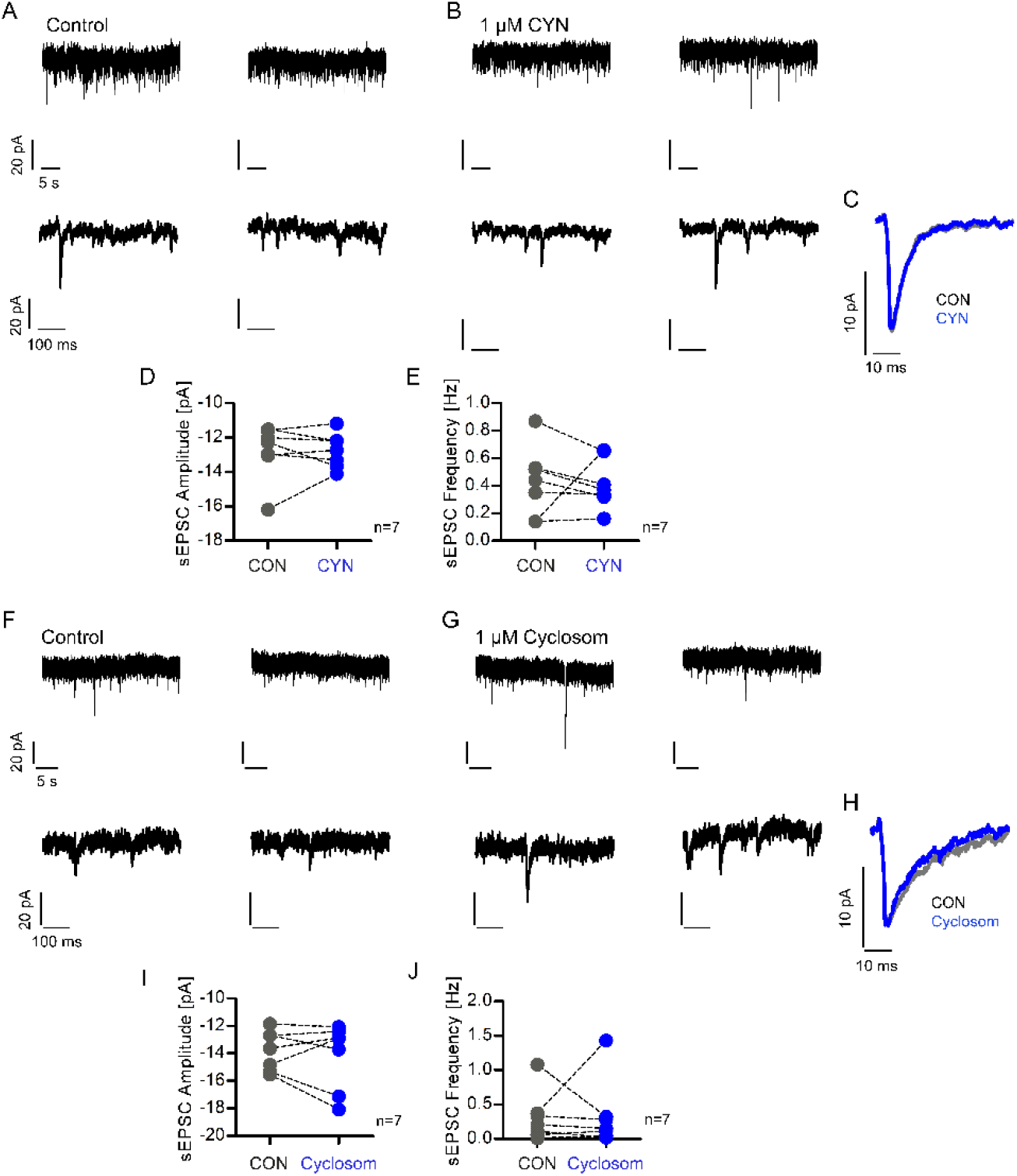
SSTR antagonism has no effect on spontaneous excitatory transmission in L2/3 PCs of the aCC. A) Upper panel: Depiction of 2 representative urrent traces of sEPSC recordings under control conditions in the presence of icuculline. Lower panel: Expanded current traces of the corresponding current traces depicted in the upper panel. B) Upper panel: Depiction of 2 representative current traces of sEPSC recordings of the same neuron as in A) after addition of CYN154806 to the bath. Lower panel: Expanded current traces of the corresponding current traces depicted in the upper panel. C) Average sEPSC trace of all events detected under control condition (light grey trace) and after addition of CYN154806 (blue trace) to the bath in the same neuron. D) As shown in the scatter dot plot, CYN154806 bath application had no effect on the sEPSC amplitude. The data were derived from 7 PCs. Mean sEPSC amplitude control: −12.80 ± 1.60 pA; SOM: −12.77 ± 1.01 pA, *p*=0.950, paired t test. E) The scatter dot plot depicts the sEPSC frequencies before and after addition of the drug. The data were derived from 7 PCs. Mean sEPSC frequency control: 0.43 ± 0.25 Hz, mean sEPSC frequency CYN: 0.42 ± 0.18 Hz, *p*=0.907, Paired t test. F) Upper panel: Depiction of 2 representative current traces of sEPSC recordings under control conditions in the presence of bicuculline. Lower panel: Expanded current traces of the corresponding current traces depicted in the upper panel. G) Upper panel: Depiction of 2 representative urrent traces of sEPSC recordings of the same neuron as in A) after addition of Cyclosomatostatin (cylcosom) to the bath. Lower panel: Expanded current traces of the corresponding current traces depicted in the upper panel. H) Average sEPSC trace of all events detected under control condition (light grey trace) and after addition of Cyclosom (blue trace) to the bath in the same neuron. I) The scatter dot plot depicts the mean sEPSC amplitudes before and after addition of the drug. The data were derived from 7 PCs. Mean sEPSC amplitude control: − 13.79 ± 1.45 pA; SOM: −14.17 ± 2.41 pA, *p*=0.538, paired t test. J) The scatter dot plot depicts the sEPSC frequencies before and after addition of the drug. The data were derived from 7 PCs. Mean sEPSC frequency control: 0.31 ± 0.37 pA; SOM: 034 ± 0.50 pA, *p*=0.895, paired t test.

**S2 Fig.**
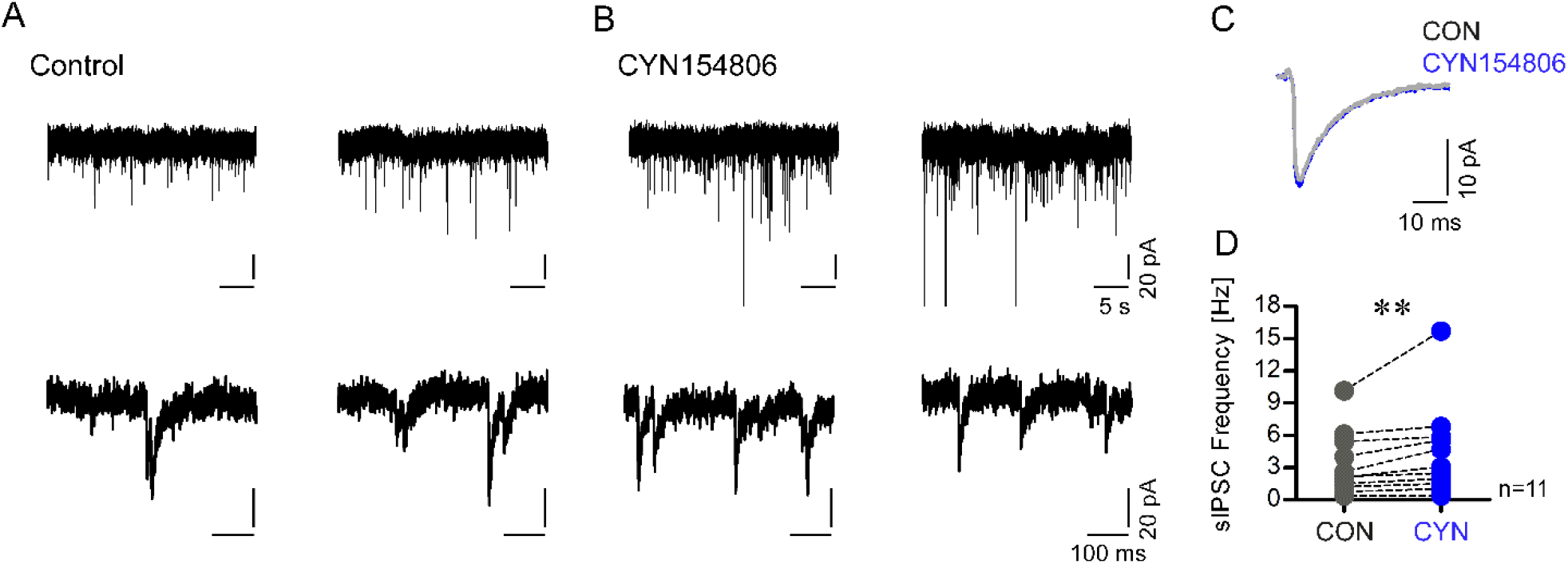
SSTR antagonism increases spontaneous inhibitory transmission in L2/3 PCs of the aCC. A) Upper panel: Depiction of 2 representativecurrent traces of sIPSC recordings under control conditions in the presence of NBQX and −AP5. Lower panel: Expanded current traces of the corresponding traces shown in the upper panel to depict individual sIPSCs. B) Upper panel: 2 representative current traces of IPSC recordings of the same neuron as in A) in the presence of CYN154806 (CYN). Lower panel: Expanded current traces of the corresponding traces in the upper panel to depict single sIPSCs. C) Average sIPSC trace of all events detected under control condition (light grey trace) and after addition of CYN (blue trace) to the bath in the same neuron. D) As shown in the dot plot, CYN bath application significantly increased the sIPSC frequency. The scatter dot plot depicts the sIPSC frequencies before and after addition of CYN. The data were derived from recordings of 11 PCs. Mean sIPSC frequency control: 3.32 ± 2.92 Hz, mean sIPSC frequency CYN: 4.45 ± 4.30 Hz, *p*=0.005, Wilcoxon signed rank test.

**S3 Fig.**
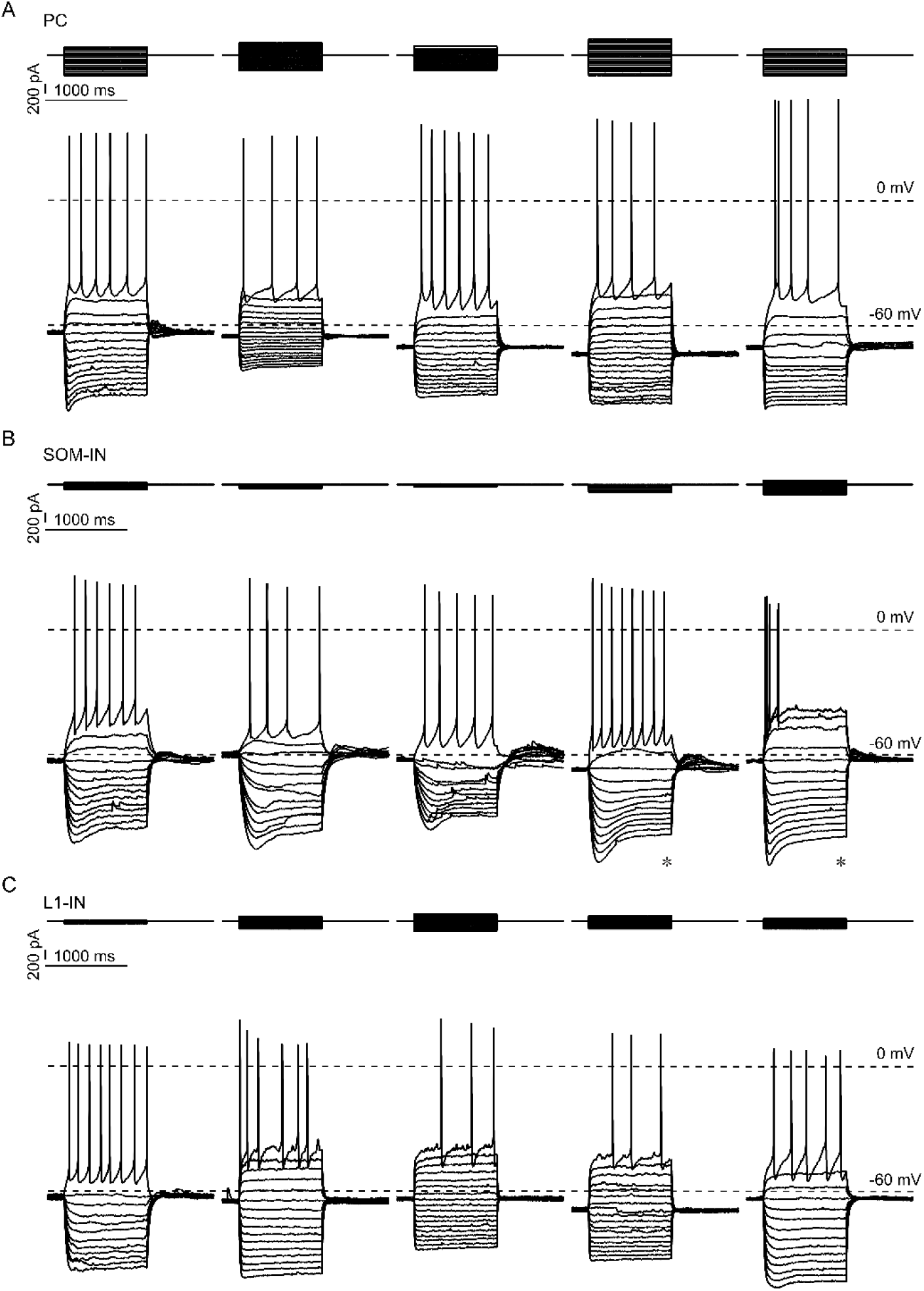
Current-Voltage relationship in L2/3 PCs, SOM-INs and L1-INs. A) Representative voltage traces (lower panel) upon injection of a series of hyper- and depolarizing current steps (upper panel) into 5 individual L2/3 PCs are depicted. B) Depiction of representative voltage traces (lower panel) of 5 individual L2/3 SOM-INs upon injection of a series of hyper- and depolarizing current steps (upper panel). Note the large sag potential (asterisk) in SOM-INs. C) Representative current- (upper panel) voltage (lower panel) traces of 5 individual L1-INs. The majority of L1-INs were late-spiking upon injection of a near threshold current and exhibited only little to no sag potential.

**S4 Fig.**
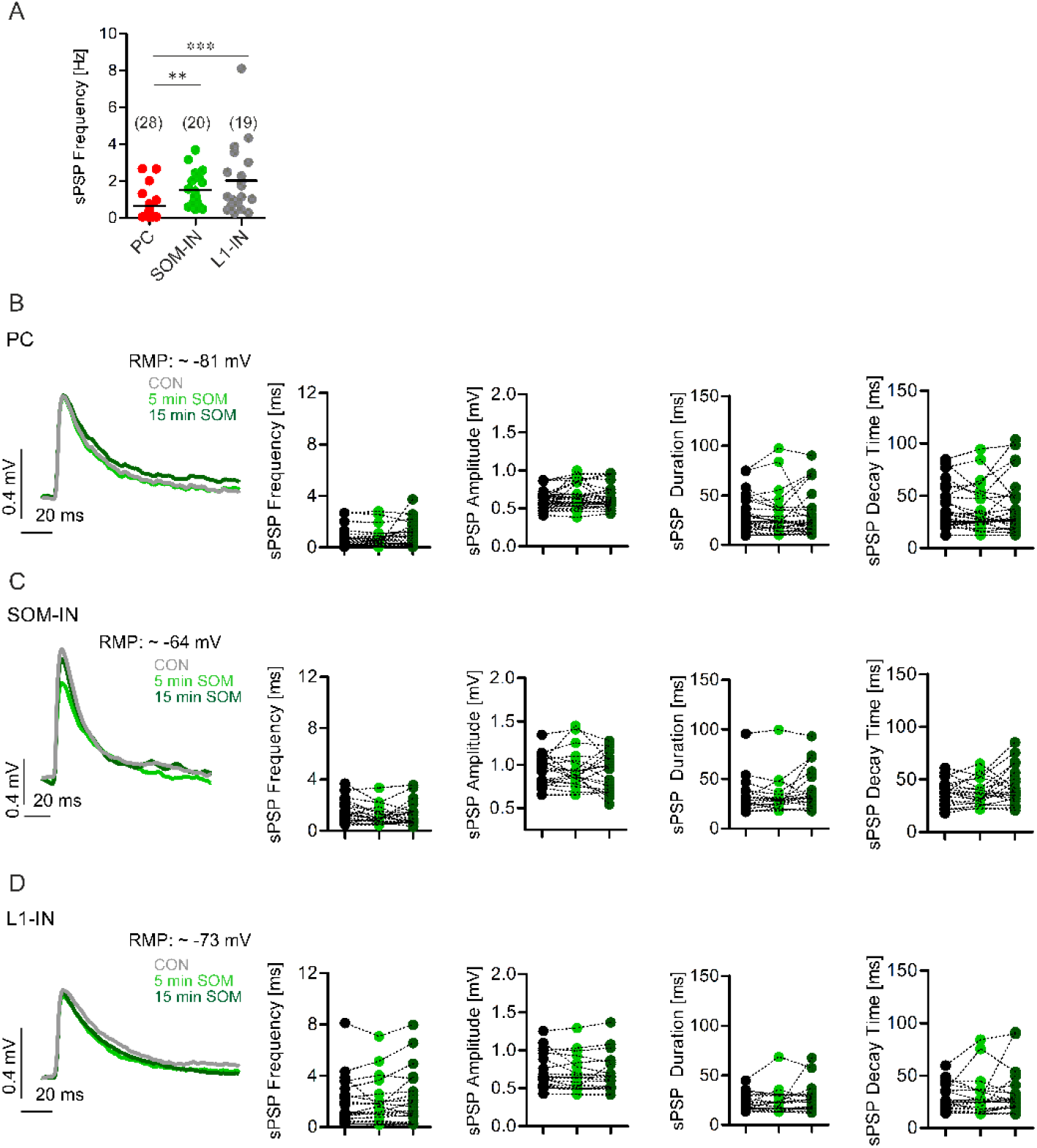
Summary of frequency and kinetics of postsynaptic potentials in L2/3 PCs, L2/3 SOM-INs and L1-INs. A) Dot plot showing the mean frequency of spontaneous postsynaptic potentials (sPSPs) recorded in PCs (red), SOM-INs (green), L1-INs (grey). L1-INs receive sPSPs at the highest, L2/3 PCs at the lowest frequency. The numbers in brackets indicate the number of cells recorded (± SEM; mean sPSP frequency L2/3 PCs: 0.65 ± 0.14 Hz; SOM-INS: 1.51 ± 0.21 Hz; L1-INs: 2.03 ± 0.44 Hz. Comparison frequency PC vs. L1-IN: p<0.001, Kruskal-Wallis test with Dunn’s multiple comparison test. Comparison frequency PC vs. SOM-IN: p<0.01, Kruskal-Wallis test with Dunn’s multiple comparison test. Comparison frequency SOM-IN vs. L1-IN: p>0.05, Kruskal-Wallis test with Dunn’s multiple comparison test). B) Left panel: Averaged sPSP under control condition (grey), and 5 (light green) and 15 (dark green) min after onset of SOM bath application taken from the same L2/3 PC. Right panel: Scatter dot plots of sPSP frequency, sPSP amplitude, sPSP duration and sPSP decay time under control condition (grey) and 5 (light green) and 15 (dark green) min after onset of SOM bath application. C) Left panel: Averaged sPSP under control condition (grey), and 5 (light green) and 15 (dark green) min after onset of SOM bath application taken from the same L2/3 SOM- IN. Right panel: Scatter dot plots of sPSP frequency, sPSP amplitude, sPSP duration and sPSP decay time under control condition (grey) and 5 (light green) and 15 (dark green) min after onset of SOM bath application. D) Left panel: Averaged sPSP under control condition (grey), and 5 (light green) and 15 (dark green) min after onset of SOM bath application taken from the same L1-IN. Right panel: Scatter dot plots of sPSP frequency, sPSP amplitude, sPSP duration and sPSP decay time under control condition (grey) and 5 (light green) and 15 (dark green) min after onset of SOM bath application.

**Fig S5.**
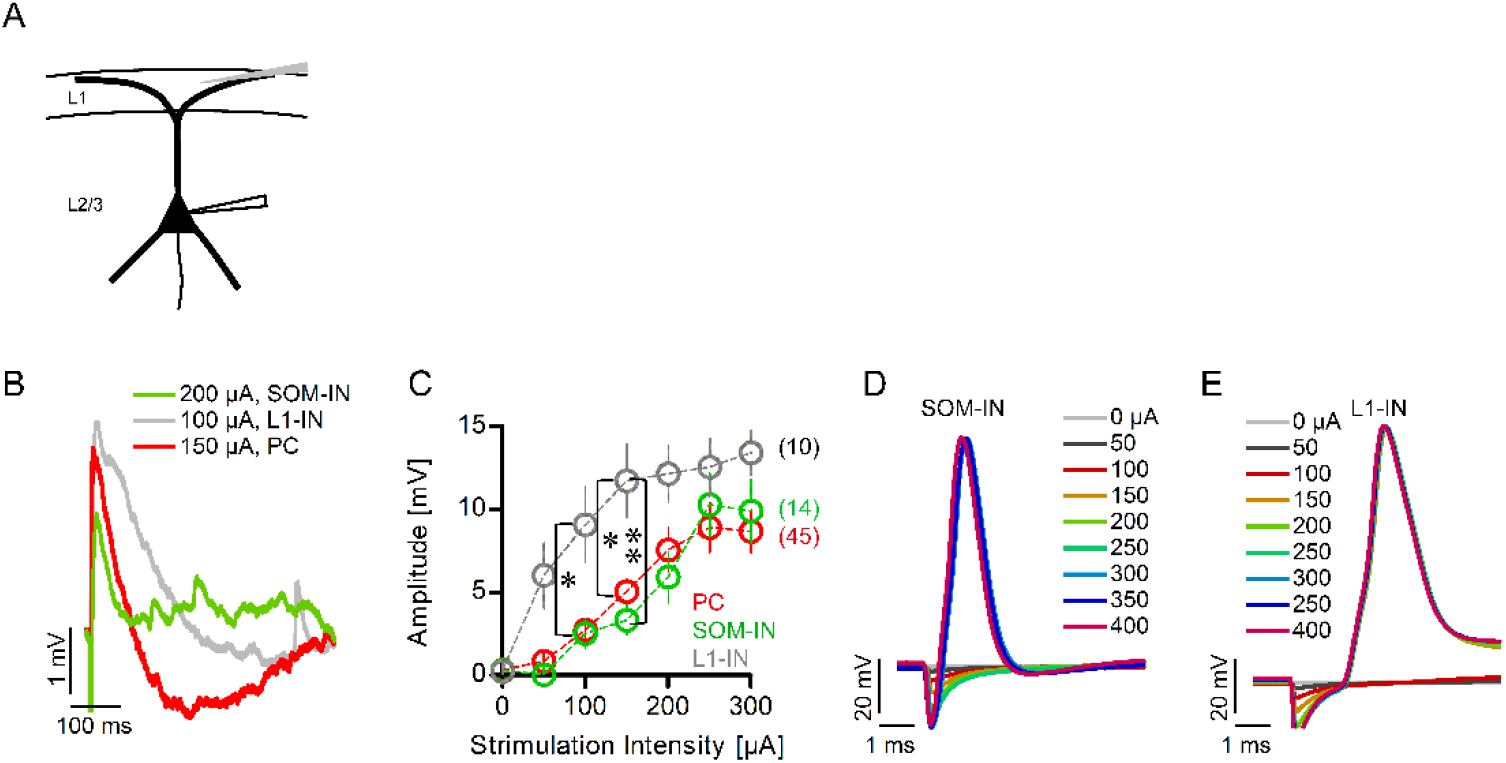
A) Scheme of the recording setup. Evoked postsynaptic potentials (ePSPs) were recorded in L2/3 PCs, SOM-INs and L1-INs after electrical stimulation in L1 at a site perpendicular to the recorded neurons. B) Merged average traces of ePSPs in a PC (red trace), a SOM-IN (green trace) and a L1-IN (grey trace). C) The amplitude of the ePSPs in PCs (red), SOM-INs (green) and L1-INs (grey) was plotted as a function of the stimulation intensity. Numbers in brackets indicate numbers of recorded cells (all data are mean ± SEM). (Amplitude of ePSPS at 0 µA: PC, 0.3 ± 0.05 mV; SOM-IN,0.28 ± 0.11 mV; L1-IN, 0.22 ± 0.07 mV. 50 µA: PC, 0.83 ± 0.17 mV; SOM-IN, 0.02 ± 0.1 mV, L1-IN, 6.03 ± 1.97 mV. 100 µA: PC, 2.71 ± 0.44 mV; SOM- IN, 2.5 ± 0.90 mV; L1-IN, 9.07 ± 2.29 mV. 150 µA: PC, 5.05 ± 0.64 mV; SOM-IN 3.31 ± 0.91 mV; L1-IN 11.73 ± 2.19 mV. 200 µA: PC, 7.56 ± 1.34 mV; SOM-IN, 5.93 ± 1.55 mV, L1-IN, 12.14 ± 1.71 mV. 250 µA: PC, 8.91 ± 1.52 mV, SOM-IN, 10.29 ± 1.94 mV; L1-IN, 12.56 ± 1.76 mV. 300 µA: PC, 8.68 ± 1.29 mV; SOM- IN 9.90 ± 1.94mV; L1-IN, 13.43 ± 1.35 mV). At 100 µA, the ePSP amplitude in L1-IN is significantly higher compared to PCs (p<0.05, Two-way ANOVA with Bonferroni post test). At 150 µA, the ePSP amplitude in L1-INs is significantly higher compared to either PCs or SOM-INs (L1-IN vs. PC: p<0.05; L1-IN vs SOM-IN: p<0.01, Two-way ANOVA with Bonferroni post test). D, E) L1 stimulation evoked antidromic spikes in SOM- INs (D) and in L1-INs (E).

